# *In vivo* recombination of *Saccharomyces eubayanus* maltose-transporter genes yields a chimeric transporter that enables maltotriose fermentation

**DOI:** 10.1101/428839

**Authors:** Nick Brouwers, Arthur R. Gorter de Vries, Marcel van den Broek, Susan M. Weening, Tom D. Elink Schuurman, Niels G. A. Kuijpers, Jack T. Pronk, Jean-Marc G. Daran

## Abstract

*Saccharomyces pastorianus* lager-brewing yeasts are aneuploid *S. cerevisiae* x *S. eubayanus* hybrids, whose genomes have been shaped by domestication in brewing-related contexts. In contrast to most *S. cerevisiae* and *S. pastorianus* strains, *S. eubayanus* cannot utilize maltotriose, a major carbohydrate in brewer’s wort. Accordingly, *S. eubayanus* CBS 12357^⊤^ harbors four *SeMALT* maltose-transporter genes, but no genes resembling the *S. cerevisiae* maltotriose-transporter gene *ScAGT1* or the *S. pastorianus* maltotriose-transporter gene *SpMTY1*. To study the evolvability of maltotriose utilization in *S. eubayanus* CBS 12357^⊤^, maltotriose-assimilating mutants obtained after UV mutagenesis were subjected to laboratory evolution in carbon-limited chemostat cultures on maltotriose-enriched wort. An evolved strain showed improved maltose and maltotriose fermentation, as well as an improved flavor profile, in 7-L fermenter experiments on industrial wort. Whole-genome sequencing revealed a novel mosaic *SeMALT413* gene, resulting from repeated gene introgressions by non-reciprocal translocation of at least three *SeMALT* genes. The predicted tertiary structure of S*e*Malt413 was comparable to the original S*e*Malt transporters, but overexpression of *SeMALT413* sufficed to enable growth on maltotriose, indicating gene neofunctionalization had occurred. The mosaic structure of *SeMALT413* resembles the structure of *S. pastorianus* maltotriose-transporter gene *SpMTY1*, which has sequences with high similarity to alternatingly *ScMALx1* and *SeMALT3*. Evolution of the maltotriose-transporter landscape in hybrid *S. pastorianus* lager-brewing strains is therefore likely to have involved mechanisms similar to those observed in the present study.

**Author Summary:** Fermentation of the wort sugar maltotriose is critical for the flavor profile obtained during beer brewing. The recently discovered yeast *Saccharomyces eubayanus* is gaining popularity as an alternative to *S. pastorianus* and *S. cerevisiae* for brewing, however it is unable to utilize maltotriose. Here, a combination of non-GMO mutagenesis and laboratory evolution of the *S. eubayanus* type strain CBS 12357^⊤^ was used to enable maltotriose fermentation in brewer’s wort. A resulting *S. eubayanus* strain showed a significantly improved brewing performance, including improved maltose and maltotriose consumption and a superior flavor profile. Whole genome sequencing identified a novel transporter gene, *SeMALT413*, which was formed by recombination between three different *SeMALT* maltose-transporter genes. Overexpression of *SeMALT413* in CBS 12357^⊤^ confirmed its neofunctionalization as a maltotriose transporter. The mosaic structure of the maltotriose transporter SpMty1 in *S. pastorianus* resembles that of S*e*Malt413, suggesting that maltotriose utilization likely emerged through similar recombination events during the domestication of current lager brewing strains.

## Introduction

*Saccharomyces eubayanus* was discovered in Patagonia and identified as the non-S. *cerevisiae* parental species of hybrid *S. pastorianus* lager-type beer brewing yeasts (1, 2). While *S. eubayanus* has only been isolated from the wild (3-5), *S. cerevisiae* is strongly associated with biotechnological processes, including dough leavening, beer brewing and wine fermentation (6). The raw material for beer brewing is wort, a complex medium containing a fermentable sugar mixture of 60% maltose, 25% maltotriose and 15% glucose (7). While most *S. cerevisiae* strains utilize all three sugars, *S. eubayanus* strains cannot utilize maltotriose (8-10). In *Saccharomyces*, the ability to utilize maltose and maltotriose is associated with *MAL* loci which are present on up to five different chromosomes (11). *MAL* loci typically harbor genes from up to three gene families: a *MALT* maltose proton-symporter gene, a *MALS* α-glucosidase gene which hydrolyses sugars into glucose, and a *MALR* regulator gene that induces the transcription of *MALT* and *MALS* genes in the presence of maltose (12). In *S. cerevisae*, most *MAL* loci harbor an *ScMalx1* transporter, which transports maltose and other disaccharides, such as turanose and sucrose (13, 14), but cannot import the trisaccharide maltotriose (15). However, the *MAL1* locus located on chromosome VII of *S. cerevisiae* contains *ScAGT1*, a transporter gene with only 57% nucleotide identity with *ScMALx1* transporter genes. *ScAGT1* encodes a broad-substrate-specificity sugar-proton symporter that enables maltotriose uptake (15-17). In *S. eubayanus* four *MAL* loci harbor a *MALT* gene with high homology to *ScMALx1* genes: *SeMALT1, SeMAL2, SeMALT3* and *SeMALT4* (18). Deletion of these genes in *S. eubayanus* type strain CBS 12357^⊤^ indicated that its growth on maltose relies on expression of *SeMALT2* and *SeMALT4* (9). *SeMALT1* and *SeMALT3* were found to be poorly expressed in the presence of maltose in this strain, supposedly due to incompleteness of the *MAL* loci which harbor them. However, no homolog of *ScAGT1* was found in the genome of CBS 12357^⊤^, and neither CBS 12357^⊤^ nor its derivatives overexpressing *SeMALT* genes were able to utilize maltotriose (9).

Laboratory-made *S. cerevisiae* x *eubayanus* hybrids combined the fermentative capacity and sugar utilization of *S. cerevisiae* with the ability of *S. eubayanus* to grow at low temperatures (8, 19, 20). Most likely, maltotriose utilization in these laboratory hybrids was enabled by the *ScAGT1* gene in the *S. cerevisiae* parental genome. Paradoxically, *S. pastorianus* strains that utilize maltotriose contain a non-functional, truncated *ScAGT1* allele (21). In such strains, maltotriose utilization has been attributed to two *S. pastorianus-specific* genes. *SpMTY1* shares 90% sequence identity with *ScMALx1* genes and enabled both maltose and maltotriose transport, with a higher affinity for the latter (22, 23). *SpMTY1* also shows sequence similarity with *SeMALT* genes (24, 25). The second gene, named *SeAGT1* because it was found in the *S. eubayanus* subgenome of *S. pastorianus* strains, shares 85% sequence identity with *ScAGT1* (26). In accordance with their sequence similarity, SeAgt1 and ScAgt1 both enable high-affinity maltotriose import (27). Despite their presence in the *S. pastorianus* genome, the maltotriose transporter genes *SpMTY1* and *SeAGT1* were not found in the genome of *S. eubayanus* CBS 12357^⊤^ (9, 18).

The *MALT* transporter genes in *S. eubayanus, S. cerevisiae* and *S. pastorianus* are localized to the subtelomeric regions (9, 14, 15, 18, 22, 23), which are gene-poor and repeat-rich sequences adjacent to the telomeres (28-30). These regions are known hotspots of genetic variation in *Saccharomyces* genomes (30-32). The presence of repeated sequences makes subtelomeric regions genetically unstable by promoting recombinations (33, 34). As a result, subtelomeric gene families are particularly diverse across different strains (32, 35, 36). In *S. cerevisiae*, subtelomeric gene families contain more genes than non-subtelomeric gene families, reflecting a higher incidence of gene duplications (35). As previously shown in *Candida albicans* submitted to long term laboratory evolution, the gene repertoire of the subtelomeric *TLO* family can be extensively altered due to ectopic recombinations between subtelomeric regions of different chromosomes, resulting in copy number expansion, in gene disappearance and in formation of new chimeric genes (37). Despite their common origin, genes within one family can have different functions, due to the accumulation of mutations (38, 39). *In silico* analysis of the sequences and functions of genes from the *MALT, MALS* and *MALR* gene families indicated functional diversification through gene duplication and mutation (35). Indeed, the presence of multiple gene copies can facilitate the emergence of advantageous mutations mainly by one of three mechanisms: (i) neofunctionalization, corresponding to the emergence of a novel function which was previously absent in the gene family (40), (ii) subfunctionalization, corresponding to the specialization of gene copies for part of the function of the parental gene (41) and (iii) altered expression due to gene dosage effects resulting from the increased copy number (42). While the different functions of *MALS* genes were assigned to subfunctionalization of the ancestral *MALS* gene (43), the maltotriose transporter gene *ScAGT1* was proposed to result from neofunctionalization within the *MALT* family (35). In general, the emergence of a large array of gene functions was attributed to subfunctionalization and neofunctionalization (35, 37, 43-47). However, current evidence for neofunctionalization within subtelomeric gene families is based on *a posteriori* analysis and rationalization of existing diversity. While in some cases the genetic process leading to neofunctionalization could be reconstructed at the molecular level (47-49), the emergence of a completely new function within a subtelomeric gene family was never observed within the timespan of an experiment to the best of our knowledge. However, the genetic diversity within *Saccharomyces MALT* transporters suggests that evolution of S*e*Malt transporters could lead to the emergence of a maltotriose transporter by neofunctionalization (35). Therefore, laboratory evolution may be sufficient to obtain maltotriose utilization in *S. eubayanus* strain CBS 12357^⊤^.

Laboratory evolution is a commonly-used non-GMO method for obtaining desired properties by prolonged growth and selection under conditions favoring cells which develop the desired phenotype (50, 51). Similarly as in Darwinian natural evolution, the conditions under which laboratory evolution is conducted shape the phenotypes acquired by evolved progeny by the process of survival of the fittest (52). In *Saccharomyces* yeasts, selectable properties include complex and diverse phenotypes such as high temperature tolerance, efficient nutrient utilization and inhibitor tolerance (53-56). Laboratory evolution was successfully applied to improve sugar utilization for arabinose, galactose, glucose and xylose (54, 57-59). In *S. pastorianus*, improved maltotriose uptake was successfully selected for in a prolonged chemostat cultivation on medium enriched with maltotriose (60). Theoretically, laboratory evolution under similar conditions could select *S. eubayanus* mutants which develop the ability to utilize maltotriose.

In this study, we submitted *S. eubayanus* strain CBS 12357^⊤^ to UV-mutagenesis and laboratory evolution in order to obtain maltotriose utilization under beer brewing conditions. While obtaining a non-GMO maltotriose-consuming *S. eubayanus* strain was a goal in itself for industrial beer brewing, we were particularly interested in the possible genetic mechanisms leading to the emergence of maltotriose utilization. Indeed, we hypothesized that the genetic plasticity of the four subtelomeric *SeMALT* genes of CBS 12357^⊤^ could facilitate the emergence of maltotriose transport by neofunctionalization. In *S. cerevisiae* the emergence of maltotriose transporter *ScAgt1* is attributed to neofunctionalization within the *MALT* gene family, and in *S. pastorianus*, the origin of the maltotriose transporter genes *SpMTY1* and *SeAGT1* remains to be elucidated. Therefore, the evolution process leading to maltotriose utilization in a strain with only maltose transporters, such as CBS 12357^⊤^, may provide insight in the emergence of maltotriose utilization in general.

## Results

### Mutagenesis and evolution enables *S. eubayanus* to utilize maltotriose

The *S. eubayanus* strain CBS 12357^⊤^ consumes maltose but not maltotriose, one of the main fermentable sugars in brewer’s wort (8). To select for maltotriose-consuming mutants, CBS 12357^⊤^ was sporulated, submitted to mild UV-mutagenesis (46% survival rate) and the mutagenized population was inoculated at 20 °C in synthetic medium containing 20 g L^−1^ maltotriose (SMMt) as sole carbon source. After two weeks, growth was observed and, after 3 weeks, the maltotriose concentration had decreased to 10.5 g L^−1^. After two subsequent transfers in fresh SMMt, 96 single cells were sorted into a microtiter YPD plate by fluorescence-activated cell sorting (FACS). The resulting single-cell cultures were transferred to a next microtiter SMMt plate, in which growth was monitored by OD_660_ measurements. The seven single-cell isolates with the highest final OD_660_ were selected and named IMS0637-IMS0643. To characterize growth on maltotriose, the strain CBS 12357^⊤^, the single-cell isolates IMS0637-IMS0643 and the maltotriose-consuming *S. pastorianus* strain CBS 1483 were grown in shake flasks on SMMt (Figure 1A and Supplementary Figure S1). After 187 h, *S. eubayanus* CBS 12357^⊤^ did not show any maltotriose consumption. Conversely, isolates IMS0637-IMS0643, all showed over 50% maltotriose consumption after 91 h (as compared to 43 h for CBS 1483). Upon reaching stationary phase, isolates IMS0637-IMS0643 had consumed 93 ± 2% of the initial maltotriose concentration, which was similar to the 92 % conversion reached by *S. pastorianus* CBS 1483. While these results indicated that the single cell isolates IMS0637-IMS0643 utilized maltotriose in synthetic medium, they did not consume maltotriose after 145 h of incubation in shake-flasks containing 3-fold diluted wort (Figure 1B). Under the same conditions, *S. pastorianus* CBS 1483 consumed 50% of the wort maltotriose after 145 h (Figure 1B).

**Figure 1:**
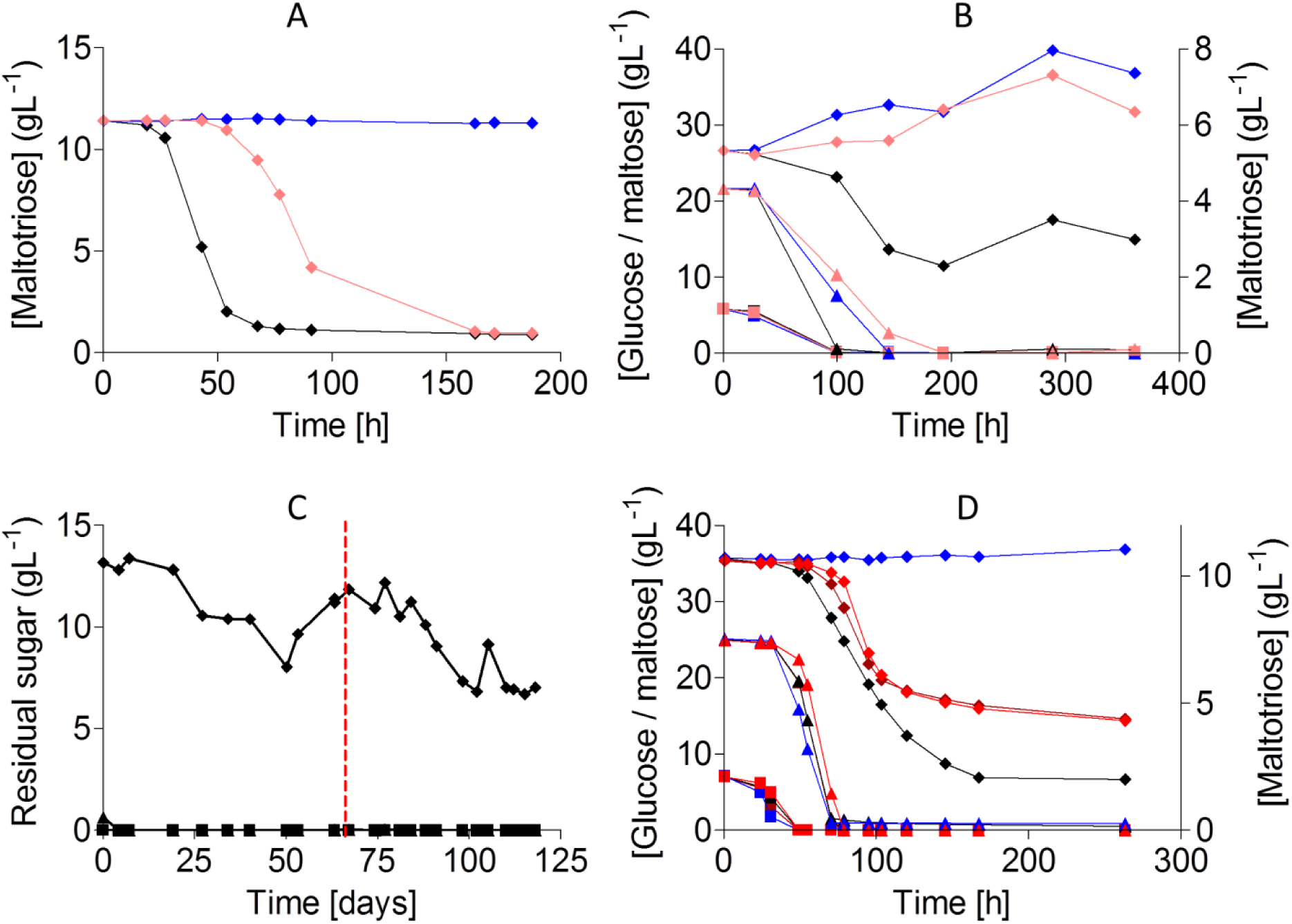
Mutagenesis and evolution to obtain maltotriose consuming *S. eubayanus*. (**A**) Characterization of *S. pastorianus* CBS 1483(black), *S. eubayanus* CBS 12357^⊤^(blue) and IMS0637 (light red) on SMMt at 20 °C. The data for IMS0637 is representative for the other mutants IMS0638-IMS0643 (Supplementary Figure S1). The average concentration of maltotriose (**♦**) and average deviation were determined from two replicates. (**B**) Characterization of *S. pastorianus* CBS 1483 (black), *S. eubayanus* CBS 12357^⊤^ (blue) and IMS0637 (light red) on wort at 20 °C. The concentrations of (**H**) glucose, (A) maltose and (**♦**) maltotriose were measured from single biological measurements. (**C**) Residual maltotriose concentration in the outflow during laboratory evolution of strains IMS0637-IMS0643 in an anaerobic chemostat at 20 °C on maltotriose enriched wort. The concentrations of (**H**) glucose, (A) maltose and (**♦**) maltotriose were measured by HPLC. The chemostat was restarted after a technical failure (red dotted line). (**D**) Characterization of *S. pastorianus* CBS 1483 (black), *S. eubayanus* CBS 12357^⊤^ (blue), IMS0750 (red) and IMS0752 (light red) on wort at 12 °C in 250 mL micro-aerobic Neubor infusion bottles. The average concentration and standard deviation of (■) glucose, (A) maltose and (♦) maltotriose were determined from three biological replicates.

Nutrient-limited growth confers a selective advantage to spontaneous mutants with a higher nutrient affinity (50, 60). Therefore, to improve maltotriose utilization under industrially relevant conditions, the pooled isolates IMS0637-IMS0643 were subjected to laboratory evolution in a chemostat culture on modified brewer’s wort. To ensure a strong selective advantage for maltotriose-consuming cells while maintaining carbon-limitation, the brewer’s wort was diluted 6-fold and complemented with 10 g L^−1^ maltotriose, yielding concentrations of 2 g L^−1^ glucose, 15 g L^−1^ maltose and 15 g L^−1^ maltotriose in the medium feed. To prevent growth limitation due to the availability of limited oxygen or nitrogen, the medium was supplemented with 10 mg L^−1^ ergosterol, 420 mg L^−1^ Tween 80 and 5 g L^−1^ ammonium sulfate (61). During the batch cultivation phase that preceded continuous chemostat cultivation, glucose and maltose were completely consumed, leaving maltotriose as the only carbon source. After initiation of continuous cultivation at a dilution rate of 0.03 h^−1^, the medium outflow initially contained 13.2 g L^−1^ of maltotriose. After 121 days of chemostat cultivation, the maltotriose concentration had progressively decreased to 7.0 g L^−1^ (Figure 1C). At that point, 10 single colony isolates were made from the culture on SMMt agar plates and incubated at 20 °C. Three single-cell lines were named IMS0750, IMS0751 and IMS0752 and selected for further characterization in micro-aerobic cultures, grown at 12 °C on 3-fold diluted wort, along with *S. eubayanus* CBS 12357^⊤^ and *S. pastorianus* CBS 1483 (Figure 1D). In these cultures, strains CBS 12357^⊤^ and IMS0751 only consumed glucose and maltose, while *S. pastorianus* CBS 1483, as well as the evolved isolates IMS0750 and IMS0752, also consumed maltotriose. After 263 h, maltotriose concentrations in cultures of strains IMS0750 and IMS0752 had decreased from 20 to 4.3 g L^−1^ maltotriose as compared to 2.0 g L^−1^ in cultures of strain CBS 1483. Due to its inability to utilize maltotriose in wort, IMS0751 was not studied further.

### Whole genome sequencing reveals a new recombined chimeric *SeMALT* gene

We sequenced the genomes of the *S. eubayanus* strain CBS 12357^⊤^, of the UV-mutagenized isolates IMS0637-IMS0643 and of the strains isolated after subsequent chemostat evolution IMS0750 and IMS0752 using paired-end Illumina sequencing. Sequencing data were mapped to a chromosome-level assembly of strain CBS 12357^⊤^ (9) to identify SNPs, INDELs and copy number changes. The genomes of the UV-mutants IMS0637, IMS0640, IMS0641 and IMS0642 shared a set of 116 SNPs, 5 INDELs and 1 copy number variation (Figure 2A, Supplementary data file 1). In addition to these shared mutations, isolates IMS0638, IMS0639 and IMS0643 carried three identical SNPs. Of the mutations in IMS0637, 34 SNPs and all 5 INDELs affected intergenic regions, 30 SNPs were synonymous, 48 SNPs resulted in amino acid substitutions and 4 SNPs resulted in premature stop codon (Supplementary data file 1). None of the 52 non-synonymous SNPs affected genes previously linked to maltotriose utilization. The only copy number variation concerned a duplication of the right subtelomeric region of CHRVIII. Read mate pairing indicated that the duplicated region was attached to the left arm of CHRII, causing the replacement of left subtelomeric region of CHRII by a non-reciprocal translocation. The affected region of CHRII harbored the *SeMALT1* gene, which is not expressed in CBS 12357^⊤^ (9).

**Figure 2:**
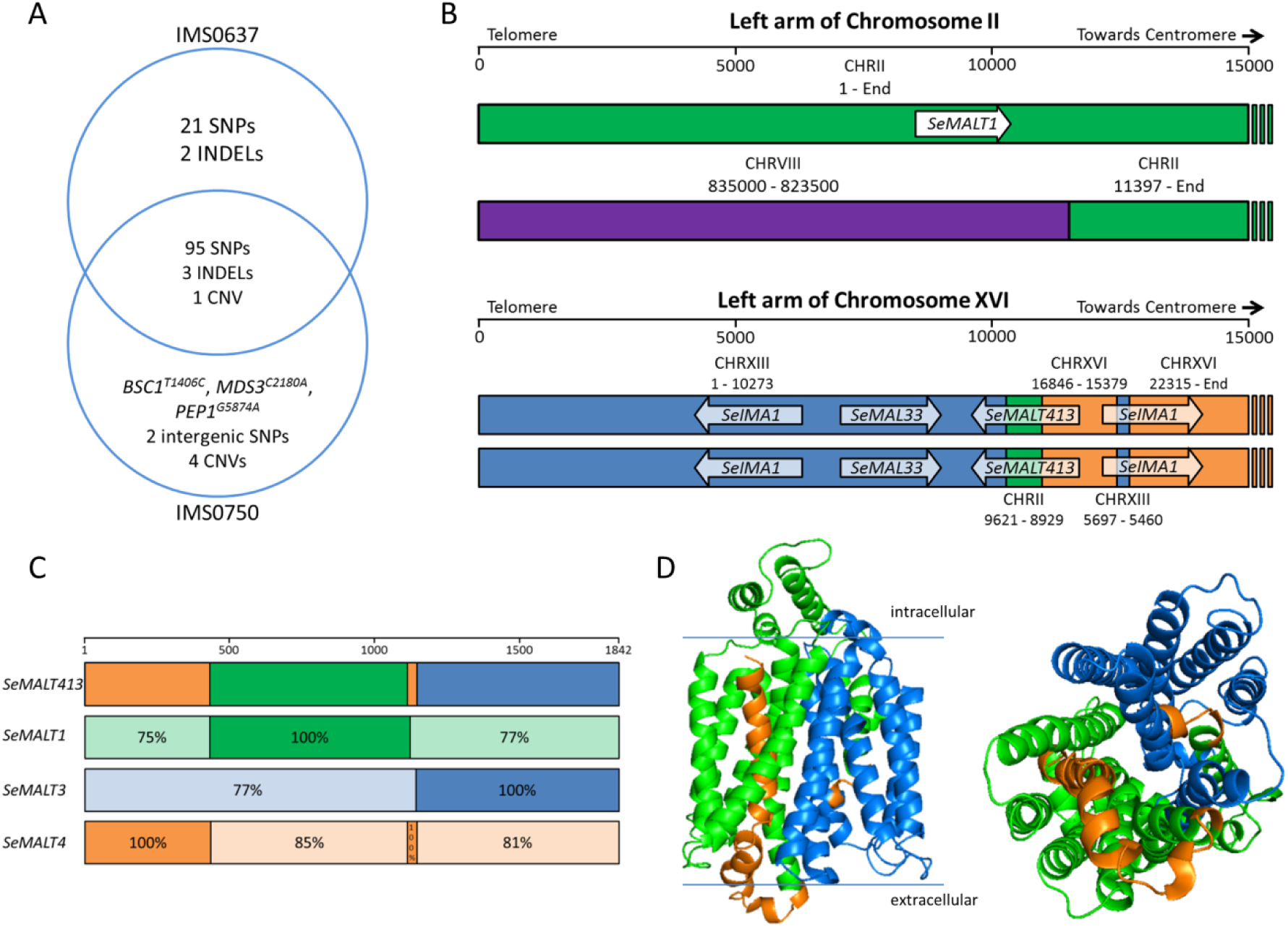
Identification of mutations in the mutagenized strain IMS0637 and the evolved strain IMS0750. (**A**) Venn diagram of the mutations found in UV-mutagenized IMS0637 and evolved IMS0750 relative to wildtype CBS 12357^⊤^. Single nucleotide polymorphisms (SNPs), small insertions and deletions (INDELs) and copy number variation (CNV) are indicated as detected by Pilon. (**B**) Recombined chromosome structures in IMS0637 and IMS0750 as detected by whole genome sequencing using MinION nanopore technology and *de novo* genome assembly. The first 15,000 nucleotides of the left arm of CHRII and CHRXVI are represented schematically. The origin of the sequence is indicated in green for CHRII, red for CHRVIII, blue for CHRXIII and orange for CHRXVI. In addition, *SeMALT* transporter genes present on the sequence are indicated by arrows. While the recombination of CHRII and CHRVIII was present in IMS0637 and IMS0750, the recombination of both copies of CHRXVI was found only in IMS0750 but not in IMS0637. The recombination on CHRXVI created the chimeric *SeMALT413* transporter gene. (**C**) Overview of the sequence similarity of the 1,842 nucleotides of *SeMALT413* relative to *SeMALT1, SeMALT3* and *SeMALT4*. The open reading frames of the genes were aligned (Supplementary Figure S2) and regions with 100% sequence identity were identified. For regions in which the sequence identity was lower than 100%, the actual sequence identity is indicated for each *SeMALT* gene. The origin of the sequence is indicated in green for CHRII, red for CHRVIII, blue for CHRXIII and orange for CHRXVI. (**D**) Prediction of the protein structure of S*e*Malt413 with on the left side a transmembrane view and on the right a transport channel view. Domains originated from *S. eubayanus* S*e*Malt transporters are indicated by the colors orange (SeMalt4 chromosome XVI), green (SeMalt1 chromosome II) and blue (SeMalt3 chromosome XIII).

Since the ability to utilize maltotriose in wort emerged only after laboratory evolution during chemostat cultivation, mutations present in the chemostat-evolved strains IMS0750 and IMS0752 were studied in more detail. With the exception of one silent SNP, IMS0750 and IMS0752 were identical and shared 100 SNPs, 3 INDELs and 5 copy number changes (Supplementary data file 1). Of these mutations, only 5 SNPs and 4 copy number changes were absent in IMS0637-IMS0643, and could therefore explain the ability to utilize maltotriose in wort (Figure 2A). The 5 SNPs consisted of two intergenic SNPs and three non-synonymous SNPs in genes with no link to maltotriose. However, the changes in copy number affected several regions harboring *SeMALT* genes: a duplication of 550 bp of CHRII including *SeMALT1* (coordinates 8,950 to 9,500), a duplication of the left arm of CHRXIII including *SeMALT3* (coordinates 1-10,275), loss of the left arm of CHRXVI (coordinates 1-15,350), and loss of 5.5 kb of CHRXVI including *SeMALT4* (coordinates 16,850-22,300). Analysis of read mate pairing indicated that the copy number variation resulted from a complex set of recombinations between chromosomes II, XIII and XVI.

The high degree of similarity of the affected *MAL* loci and their localization in the subtelomeric regions made exact reconstruction of the mutations difficult. Therefore, IMS0637 and IMS0750 were sequenced using long-read sequencing on ONT’s MinION platform, and a *de novo* genome assembly was made for each strain. Comparison of the resulting assemblies to the chromosome-level assembly of CBS 12357^⊤^ indicated that two recombinations had occurred. Both in IMS0637 and IMS0750, an additional copy of the terminal 11.5 kbp of the right arm of chromosome VIII had replaced the terminal 11.4 kbp of one of the two copies of the left arm of chromosome II (Figure 2B). This recombination was consistent with the copy number changes of the affected regions in IMS0637-IMS0643, IMS0750 and IMS0752 and resulted in the loss of one copy of the *MAL* locus harboring *SeMALT1*. In addition, the genome assembly of IMS0750 indicated the replacement of both copies of the first 22.3 kbp of CHRXVI by complexly rearranged sequences from CHRII, CHRXVIII and CHRXVI. The recombined region comprised the terminal 10,273 nucleotides of the left arm of CHRIII, followed by 693 nucleotides from CHRII, 1,468 nucleotides from CHRXVI and 237 nucleotides from CHRXIII (Figure 2B). The recombinations were non reciprocal, as the regions present on the recombined chromosome showed increased sequencing coverage while surrounding regions were unaltered. This recombination resulted in the loss of the canonical *MAL* locus harboring *SeMALT4* on chromosome XVI. However, the recombined sequence contained a chimeric open reading frame consisting of the 5’ part of *SeMALT4* from CHRXVI, the middle of *SeMALT1* from CHRII and the 3’ part of *SeMALT3* from CHRXIII (Figure 2C, Supplementary Figure S2). To verify this recombination, the ORF was PCR amplified using primers binding on the promotor of *SeMALT4* and the terminator of *SeMALT3*, yielding a fragment for strain IMS0750, but not for CBS 12357^⊤^. Sanger sequencing of the fragment amplified from strain IMS0750 confirmed the chimeric organization of the ORF, which we named *SeMALT413*. The sequence of *SeMALT413* showed 100% identity to *SeMALT4* for nucleotides 1-434 and 1113-1145, 100% similarity to *SeMALT1* for nucleotides 430-1122 and 100% similarity to *SeMALT3* for nucleotides 1141-1842 (Figure 2C). Nucleotides 1123-1140, which showed only 72% identity with *SeMALT1* and 61% identity with *SeMALT3*, were found to represent an additional introgression (Figure 2B). While the first 434 nucleotides can be unequivocally attributed to *SeMALT4* due to a nucleotide difference with *SeMALT2*, the nucleotides 1123-1140 are identical in *SeMALT2* and *SeMALT4*. Therefore, this part of the sequence of *SeMALT413* might have come from *SeMALT2* on CHRV or from *SeMALT4* on CHRXVI. Overall, *SeMALT413* showed a sequence identity of only 85 to 87% with the original *SeMALT* genes, with the corresponding protein sequence exhibiting between 52 and 88% identity. We therefore hypothesized that the recombined S*e*Malt413 transporter might have an altered substrate specificity and thereby enable maltotriose utilization.

The tertiary structure of the chimeric *SeMALT413* gene was predicted with SWISS-MODEL (https://swissmodel.expasy.org/), based on structural homology with the *Escherichia coli* xylose-proton symporter XylE (62), which has previously been used as a reference to model the structure of ScAgt1 (63). Similarly to the maltose transporters in *Saccharomyces*, XylE is a proton symporter belonging to the major facilitator superfamily with a transmembrane domain composed of 12 α-helixes (Supplementary Figure S3). The same structure was predicted for S*e*Malt413, with 1 α-helix formed exclusively by residues from S*e*Malt4, 4 α-helixes formed by residues from S*e*Malt1 and 5 α-helixes formed exclusively by residues from S*e*Malt3 (Figure 2D). The remaining two a-helixes were composed of residues from more than one transporter. Since the first 100 amino acids were excluded from the model due to absence of similar residues in the xylose symporter reference model, the structure prediction underestimated the contribution of *Se*Malt4. The three-dimensional arrangement of the a-helixes of S*e*Malt413 was almost identical to S*e*Malt1, S*e*Malt3 and S*e*Malt4, indicating that it retained the general structure of a functional maltose transporter (Supplementary Figure S4).

### Introduction of the *SeMALT413* gene in wildtype CBS 12357^⊤^ enables maltotriose utilization

The small structural differences identified between S*e*Malt413 and the wild-type *S. eubayanus* Malt transporters could not be used to predict the ability of S*e*Malt413 to transport maltotriose (63). Therefore, to investigate its role in maltotriose transport, *SeMALT413* and, as a control, *SeMALT2* were overexpressed in the wild-type strain *S. eubayanus* CBS 12357^⊤^ (Figure 3A and Supplementary Figure S5). Growth of the resulting strains *S. eubayanus* IMX1941 (*SeSGA1*Δ*::ScTEF1_pr_-SeMALT2-*ScCYC1_ter_) and IMX1942 (*SeSGA1*Δ::*ScTEF1*_pr_-*SeMALT413*-*ScCYC1*_ter_), as well as the wild-type strain CBS 12357^⊤^ and the evolved isolate IMS0750 was tested on SM supplemented with different carbon sources (Supplementary Figure S6). On glucose, strains IMX1941 and IMX1942 exhibited the same specific growth rate of 0.25 ± 0.01 h^−1^ as CBS 12357^⊤^, while IMS0750 grew faster with a growth rate of 0.28 ± 0.01 h^−1^. Glucose was completely consumed after 33 h (Figure 3B). On maltose, the specific growth rates of CBS 12357^⊤^, IMX1941, IMX1942 and IMS0750 ranged between 0.17 and 0.19 h^−1^ and did not differ significantly. Maltose was completely consumed after 43 h (Figure 3C). On maltotriose, only the evolved mutant IMS0750 and reverse engineered strain IMX1942 (*ScTEF1*_pr_*-SeMALT413-ScCYC1*_ter_) showed growth. IMS0750 grew with a specific growth rate of 0.19 ± 0.01 h^−1^ and consumed 55% of maltotriose within 172 h. Over the same period, IMX1942 grew at 0.03 ± 0.00 h^−1^ and consumed 45% of the maltotriose after 172 h (Figure 3D), demonstrating the capacity of *SeMALT413* to transport maltotriose.

**Figure 3:**
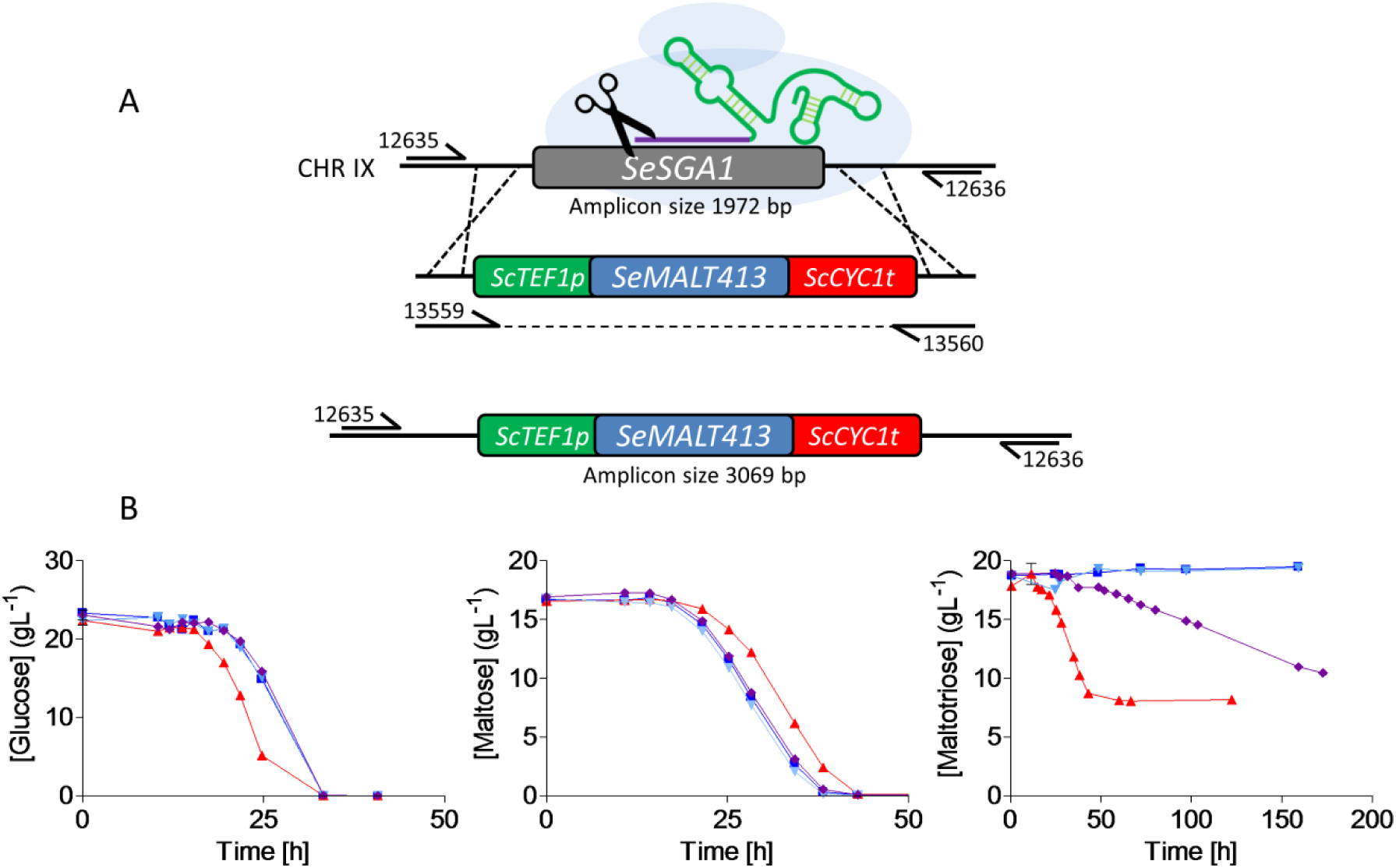
Reverse engineering of *SeMALT413* in CBS 12357^⊤^ and characterization of transporter functionality in SM. (**A**) Representation of the CRISPR-Cas9 gRNA complex (after self-cleavage of the 5’ hammerhead ribozyme and a 3’ hepatitis-δ virus ribozyme from the expressed gRNA) bound to the *SeSGA1* locus in CBS 12357^⊤^. Repair fragment with transporter cassette *ScTEF1p-SeMALT413-ScCYC1t* was amplified from pUD814(SeMALT413) with primers 13559/13560 and contains overhangs with the *SeSGA1* locus for recombination. *SeSGA1* was replaced by the *ScTEF1p-SeMALT413-ScCYC1t* cassette. Correct transformants were checked using primers 12635/12636 upstream and downstream of the *SeSGA1* locus (Supplementary Figure S5). Strains were validated using Sanger sequencing. (**B**) Characterization of (■) CBS 12357, (▲) IMS0750, (▼) IMX1941, (♦) IMX1942 on SM glucose, maltose and maltotriose. Strains were cultivated at 20 °C and culture supernatant was measured by HPLC. Data represent average and standard deviation of three biological replicates.

### The *SpMTY1* maltotriose transporter gene displays a similar chimeric structure as *SeMALT413*

The mosaic structure of the maltotriose transporter gene *SeMALT413* led us to reinvestigate the sequence of maltotriose transporters in *Saccharomyces* genomes. The sequence similarity of *ScAGT1* and *SeAGT1* to maltose transporters from the *MALT* family such as *ScMAL31* is roughly homogenous over their coding region. In contrast, the identity of some segments of *SpMTY1* relative to *ScMAL31* deviates strongly from the average identity of 89% (22). Indeed, sequence identity with *ScMAL31* of *S. cerevisiae* S288C (64) is above 98% for nucleotides 1-439, 627-776, 796-845, 860-968 and 1,640-1,844, while it is only 79% for nucleotides 440-626, 65% for nucleotides 777-795, 50% for nucleotides 846-859 and 82% for nucleotides 969-1,639 (Supplementary Figure S7). Alignment of the sequences of *S. eubayanus* CBS 12357^⊤^ *SeMALT* genes (9) to *SpMTY1* showed high sequence identity with *SeMALT3* across several regions that showed significant divergence from the corresponding *ScMAL31* sequences: 91% similarity for nucleotides 478-533, 94% similarity for nucleotides 577-626 and 94% similarity for nucleotides 778-794 (Supplementary Figure S7). These observations would indicate that the evolution of *SpMTY1* might have involved introgression events similar to those responsible for the *SeMALT413* neofunctionalization described in the present study. However, introgressions from *SeMALT* genes cannot explain the entire *SpMTY1* gene structure. Its evolution may therefore have involved multiple introgressions, similarly as for *SeMALT413*. While most regions with low similarity to *ScMAL31* and *SeMALT3* were too short to identify their provenance, the sequence corresponding to the 969^th^ to 1,639^th^ nucleotide of *SpMTY1* could be blasted on NCBI. In the S288C genome, *ScMAL31* was the closest hit with 82% identity. However, when blasting the sequence against the full repository excluding *S. pastorianus* genomes, the closest hit was the orthologue of *ScMAL31* on chromosome VII of *S. paradoxus* strain YPS138. In addition to an 89% similarity to nucleotides 969-1,639 of *SpMTY1, SparMAL31* had a similarity of 94% for nucleotides 544-575 and of 93% for nucleotides 846-859 (Supplementary Figure S7). Therefore, *SparMAL31* may have contributed sequence to the 3’ part of *SpMTY1* by horizontal gene transfer.

### Applicability of a maltotriose-consuming *S. eubayanus* strain for lager beer brewing

*S. eubayanus* strains are currently used for industrial lager beer brewing (9). To test the evolved strain IMS0750 under laboratory-scale brewing conditions, its performance was compared with that of its parental strain CBS 12357^⊤^ in 7-L cultures grown on high-gravity (16.6 ° Plato) wort (Figure 4). After 333 h, IMS0750 had completely consumed all glucose and maltose, and the concentration of maltotriose had dropped from 19.3 to 4.7 g L^−1^ (Figure 4). In contrast, CBS 12357^⊤^ did not utilize any maltotriose. In addition to its improved maltotriose utilization, IMS0750 also showed improved maltose consumption: maltose was completely consumed within 200 h, while complete maltose consumption by strain CBS 12357^⊤^ took 333 h (Figure 4). Consistent with its improved sugar utilization, the final ethanol concentration in cultures of strain IMS0750 was 18.5% higher than in corresponding cultures of strain CBS 12357^⊤^ (Figure 4). Brewing-related characteristics of IMS0750 were further explored by analyzing production of aroma-defining esters, higher alcohols and diacetyl. Final concentrations of esters and higher alcohols were not significantly different in cultures of the two strains, with the exception of isoamylacetate, which showed a 240 % higher concentration in strain IMS0750 (Table 1). In addition, while the concentration of the off-flavour diacetyl remained above its taste threshold of 25 μg L^−1^ after 333h for CBS 12357^⊤^, it dropped below 10 μg L^−1^ for IMS0750 (Table 1).

**Figure 4:**
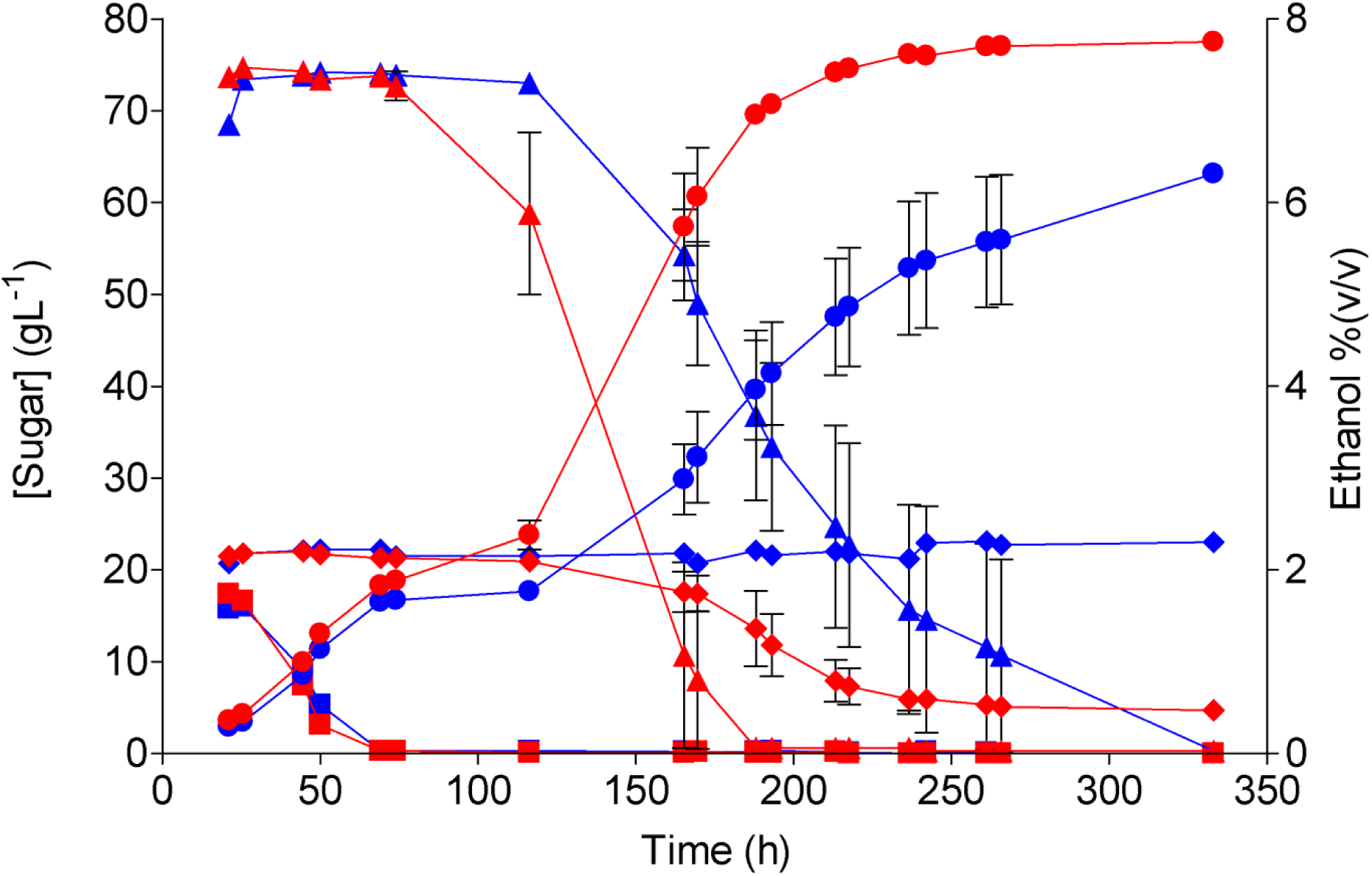
Extracellular metabolite profiles of *S. eubayanus* strains CBS 12357^⊤^ and IMS0750 in high-gravity wort at 7-L pilot scale. Fermentations were performed on wort with a gravity of 16.6 °Plato. The average concentrations of glucose (■), maltose (▲), maltotriose (♦) and ethanol (●) are shown for duplicate fermentations of CBS 12357^⊤^ (blue) and IMS0750 (red). The average deviations are indicated.

**Table 1:** Concentrations of alcohols, esters and diacetyl after fermentation of wort with a gravity of 16.6 °P by S. eubayanus strains CBS 12357^⊤^ and IMS0750. The data correspond to the last time point (330 h) of the fermentations shown in Figure 4. The average and average deviation of duplicate fermentations are shown for each strain.

| Compound | Unit | CBS 12357<sup>T</sup> | IMS0750 |
| --- | --- | --- | --- |
| Methanol | mg L<sup>-1</sup> | 3.3 ± 0.3 | 3.7 ± 0.3 |
| Propanol | mg L<sup>-1</sup> | 23.7 ± 2.1 | 24.1 ± 0.9 |
| Isobutanol | mg L<sup>-1</sup> | 48.5 ± 2.4 | 42.9 ± 7.2 |
| Amyl alcohol | mg L<sup>-1</sup> | 138.5 ± 9.0 | 155.9 ± 6.4 |
| Diacetyl | μg L<sup>-1</sup> | 43.8 ± 22.9 | 7.5 ± 0.2 |
| Ethylacetate | mg L<sup>-1</sup> | 24.5 ± 5.5 | 26.1 ± 0.8 |
| Isoamylacetate | mg L<sup>-1</sup> | 1.4 ± 0.6 | 3.1 ± 0.3 |

## Discussion

UV mutagenesis and subsequent laboratory evolution in maltotriose-limited chemostat cultures yielded *S. eubayanus* strains that were able to ferment maltotriose in laboratory-scale wort fermentation experiments. Whole genome sequencing of the mutants before and after the emergence of maltotriose utilization in wort resulted in the identification of several recombinations affecting subtelomeric regions. All four maltose transporter genes in *S. eubayanus* CBS 12357^⊤^ are localized in subtelomeric *MAL* loci: *SeMALT1* on chromosome II, *SeMALT2* on chromosome V, *SeMALT3* on chromosome XIII and *SeMALT4* on chromosome XVI (9, 18). In the evolved strain IMS0750, a complex recombination between the subtelomeric regions of chromosomes II, XIII and XVI involved at least three of these *MAL* loci. Long-read nanopore sequencing enabled complete reconstruction of the recombined left arm of chromosome XVI, revealing recombinations between the ORFs of at least *SeMALT1, SeMALT3* and *SeMALT4*. These recombinations occurred within the open reading frame of *SeMALT4* and the newly-formed chimeric ORF *SeMALT413* encoded a full length protein with a structure comparable to that of *SeMalt* transporters. In contrast to the original *SeMALT* genes, overexpression of *SeMALT413* enabled growth on maltotriose, indicating that S*e*Malt413 acquired the ability to import maltotriose. While the emergence of a new ORF by recombination has been observed previously between the *TLO* genes of *C. albicans*, it was not associated with a new gene function (37). In contrast, the emergence of *SeMALT413* is an example of gene neofunctionalization, which occurred by recombination within genes of the subtelomeric *MALT* family.

Neofunctionalization by *in vivo* formation of chimeric sequences is reminiscent of the *in vitro* protein engineering strategy known as gene shuffling or gene fusion (65, 66). Gene shuffling involves randomized assembly of diverse DNA sequences into chimeric genes, followed by screening for novel or improved functions. Analogously to *in vitro* gene shuffling, the complex protein remodeling caused by *in vivo* formation of chimeric sequences may be particularly potent for protein neofunctionalization (67). The demonstration of neofunctionalization of a sugar transporter in *S. eubayanus* by *in vivo* gene shuffling supports the notion that gene fusion is an essential driver of evolution by accelerating the emergence of new enzymatic functions (68). Moreover, analysis of the *SpMTY1* maltotriose transporter gene revealed a chimeric structure similar to that of *SeMALT413*, albeit with alternating sequence identity with *ScMAL31, SeMALT* and *SparMAL31*. While sequences from *S. cerevisiae* and *S. eubayanus* were already present in the genome of *S. pastorianus*, the presence of sequences from *S. paradoxus* is plausible as introgressions from *S. paradoxus* were commonly found in a wide array of *S. cerevisiae* strains (31). particularly Therefore, the sequence of *SpMTY1* could have resulted from *in vivo* gene shuffling between genes from the *MALT* family, followed by accumulation of mutations. The emergence of *SeMALT413* could therefore be representative of the emergence of maltotriose utilization during the evolution of *S. pastorianus*.

No evidence of reciprocal translocations between *SeMALT1, SeMALT3* and *SeMALT4* was found in the genome of IMS0750, indicating genetic introgression via non-conservative recombinations. Such introgressions can occur during repair of double strand breaks by strand invasion of a homologous sequence provided by another chromosome and resection (69), leading to localized gene conversion and loss of heterozygosity. This model, which was proposed to explain local loss of heterozygosity of two orthologous genes in an *S. cerevisiae* x *S. uvarum* hybrid (69), provides a plausible explanation of the emergence of *SeMALT413* through non-reciprocal recombination between paralogous *SeMALT* genes in *S. eubayanus*. The mosaic sequence composition of the resulting transporter gene suggests that neofunctionalization required multiple successive introgression events. As a result of these genetic introgressions, the *SeMALT4* gene was lost. The fact that IMS0750 harbored two copies of *SeMALT413* and no copy of *SeMALT4* indicates a duplication of the newly-formed ORF at the expense of *SeMALT4* via loss of heterozygosity. As functional-redundancy enables the accumulation of mutation without losing original functions (35, 37, 38, 70), the loss of *SeMALT4* was likely facilitated by the presence of the functionally-redundant maltose transporter *SeMALT2* (9). The observation that introgressions were only found at *SeMALT4* may be due to the low number of tested mutants. However, it should be noted that introgressions in the *SeMALT1* and *SeMALT3* ORF’s would have been unlikely to be beneficial, since these genes are not expressed in CBS 12357^⊤^ (9).

This study illustrates the role of the rapid evolution of subtelomeric genes in adaptation to environmental changes. In addition, the newly-acquired ability of *SeMALT413* to transport maltotriose constitutes an example of evolution by gene neofunctionalization in the laboratory environment. The emergence of new functions is critical for the process of evolution. *A posteriori* analysis of existing gene families has provided insights on their evolutionary history and on the emergence of new functions. For example, the α-glucosidase genes from the *MALS* family emerged by expansion of an ancestral pre-duplication gene with maltose-hydrolase activity and trace isomaltose-hydrolase activity. The evolution of MALS isomaltase genes from this ancestral gene is an example of subfunctionalization: the divergent evolution of two gene copies culminating in their specialization for distinct functions which were previously present to a lesser extent in the ancestral gene. The generation of functional redundancy by gene duplication is critical to this process as it enables mutations to occur which result in loss of the original gene function without engendering a selective disadvantage (35, 37, 38, 40, 41, 70). In contrast to subfunctionalization, neofunctionalization consists of the emergence of a function which was completely absent in the ancestral gene (45). While the emergence of many genes from a large array of organisms has been ascribed to subfunctionalization and to neofunctionalization, these conclusions were based on *a posteriori* analysis of processes which had already occurred, and not on their experimental observation (35, 37, 43-47). Here we present clear experimental evidence of neofunctionalization within a laboratory evolution experiment. Furthermore, while *ex-vivo* engineering of the subtelomeric *FLO* genes had already shown that recombinations within subtelomeric gene families can alter their function (44), the ability of *SeMALT413* to transport maltotriose proves that such *in vivo* gene shuffling is relevant for evolutionary biology.

While the introduction of *SeMALT413* in CBS 12357^⊤^ via genetic engineering demonstrated its neofunctionalization, the use of GMO-strains is precluded in the brewing industry by customer acceptance issues (71). However, the non-GMO evolved *S. eubayanus* isolate IMS0750 could be tested on industrial brewing wort at 7 L scale. In addition to near-complete maltotriose conversion, the maltose consumption, isoamylacetate production and diacetyl degradation of IMS0750 were superior to CBS 12357^⊤^. Efficient maltose and maltotriose consumption, as well as the concomitantly increased ethanol production, are important factors determining the economic profitability of beer brewing processes (72). In addition, low residual sugar concentration, low concentrations of diacetyl and high concentrations of Isoamylacetate are desirable for the flavor profile of beer (73, 74). In terms of application, the laboratory evolution approach for conferring maltotriose utilization into *S. eubayanus* presented in this paper is highly relevant in view of the recent introduction of this species in industrial-scale brewing processes (9). The ability to ferment maltotriose can be introduced into other natural isolates of *S. eubayanus*, either by laboratory evolution or by crossing with evolved strains such as *S. eubayanus* IMS0750. Besides their direct application for brewing, maltotriose-consuming *S. eubayanus* strains are of value for the generation of laboratory-made hybrid *Saccharomyces* strains for brewing and other industrial applications (8, 75-77).

## Materials and methods

### Strains and maintenance

All yeast strains used and generated in this study are listed in Table 2. *S. eubayanus* type strain CBS 12357^⊤^ (1) and *S. pastorianus* strain CBS 1483 (60, 78) were obtained from the Westerdijk Fungal Biodiversity Institute (Utrecht, the Netherlands). Stock cultures were grown in YPD, containing 10 g L^−1^ yeast extract, 20 g L^−1^ peptone and 20 g L^−1^ glucose, at 20 °C until late exponential phase, complemented with sterile glycerol to a final concentration of 30% (v/v) and stored at −80 °C until further use.

**Table 2:** *Saccharomyces* strains used during this study.

| Name | Species | Relevant genotype | Origin |
| --- | --- | --- | --- |
| CBS 12357 | *S. eubayanus* | Wildtype diploid | (1) |
| IMS0637 | *S. eubayanus* | Evolved strain derived from CBS 12357 | This study |
| IMS0638 | *S. eubayanus* | Evolved strain derived from CBS 12357 | This study |
| IMS0639 | *S. eubayanus* | Evolved strain derived from CBS 12357 | This study |
| IMS0640 | *S. eubayanus* | Evolved strain derived from CBS 12357 | This study |
| IMS0641 | *S. eubayanus* | Evolved strain derived from CBS 12357 | This study |
| IMS0642 | *S. eubayanus* | Evolved strain derived from CBS 12357 | This study |
| IMS0643 | *S. eubayanus* | Evolved strain derived from CBS 12357 | This study |
| IMS0750 | *S. eubayanus* | Evolved strain derived from CBS 12357 | This study |
| IMS0751 | *S. eubayanus* | Evolved strain derived from CBS 12357 | This study |
| IMS0752 | *S. eubayanus* | Evolved strain derived from CBS 12357 | This study |
| IMX1941 | *S. eubayanus* | ΔSesga1::ScTEF1p-SeMALT2-ScCYC1t | This study |
| IMX1942 | *S. eubayanus* | ΔSesga1::ScTEF1p-SeMALT413-ScCYC1t | This study |
| CBS 1483 | *S. pastorianus* | Group II brewer's yeast, Brewery Heineken, bottom yeast, July 1927 | (78) |

### Media and cultivation

Plasmids were propagated overnight in *Escherichia coli* XL1-Blue cells in 10 mL LB medium containing 10 g L^−1^ peptone, 5 g L^−1^ Bacto Yeast extract, 5 g L^−1^ NaCl and 100 mg L^−1^ ampicillin at 37 °C. Synthetic medium (SM) contained 3.0 g L^−1^ KH_2_PO_4_, 5.0 g L^−1^ (NH_4_)_2_SO_4_, 0.5 g L^−1^ MgSO_4_, 7 H_2_O, 1 mL L^−1^ trace element solution, and 1 mL L^−1^ vitamin solution (61), and was supplemented with 20 g L^−1^ glucose (SMG), maltose (SMM) or maltotriose (SMMt) by addition of autoclaved 50% w/v sugar solutions. Maltotriose (95.8% purity) was obtained from Glentham Life Sciences, Corsham, United Kingdom. Industrial wort was provided by HEINEKEN Supply Chain B.V., Zoeterwoude, the Netherlands. The wort was supplemented with 1.5 mg L^−1^ of Zn^2+^ by addition of ZnSO_4_·7H_2_O, autoclaved for 30 min at 121°C and filtered using Nalgene 0.2 μm SFCA bottle top filters (Thermo Scientific, Waltham, MA) prior to use. Where indicated, filtered wort was diluted with sterile demineralized water. Solid media were supplemented with 20 g L^−1^ of Bacto agar (Becton Dickinson, Breda, The Netherlands). *S. eubayanus* strains transformed with plasmids pUDP052 (gRNA*_SeSGA1_*) were selected on medium in which (NH_4_)_2_SO_4_ was replaced by 5 g L^−1^ K_2_SO_4_ and 10 mM acetamide (SM_Ace_G: SMG) (79).

### Shake-flask cultivation

Shake-flask cultures were grown in 500 mL shake flasks containing 100 mL medium and inoculated from stationary-phase aerobic precultures to an initial OD_660_ of 0.1. Inocula for growth experiments on SMMt were grown on SMM. In other cases, media for growth experiments and inoculum preparation were the same. Shake flasks were incubated at 20 °C and 200 RPM in a New Brunswick Innova43/43R shaker (Eppendorf Nederland B.V., Nijmegen, The Netherlands). Samples were taken at regular intervals to determine OD_660_ and extracellular metabolite concentrations.

### Microaerobic growth experiments

Microaerobic cultivation was performed in 250 mL airlock-capped Neubor infusion bottles (38 mm neck, Dijkstra, Lelystad, Netherlands) containing 200 mL 3-fold diluted wort supplemented with 0.4 mL L^−1^ Pluronic antifoam (Sigma-Aldrich). Bottle caps were equipped with a 0.5 mm × 16 mm Microlance needle (BD Biosciences) sealed with cotton to prevent pressure build-up. Sampling was performed aseptically with 3.5 mL syringes using a 0.8 mm × 50 mm Microlance needle (BD Biosciences). Microaerobic cultures were inoculated at an OD_660_ of 0.1 from stationary-phase precultures in 50 mL Bio-One Cellstar Cellreactor tubes (Sigma-Aldrich) containing 30 mL of the same medium, grown for 4 days at 12 °C. Bottles were incubated at 12 °C and shaken at 200 RPM in a New Brunswick Innova43/43R shaker. At regular intervals, 3.5 mL samples were collected in 24 deep-well plates (EnzyScreen BV, Heemstede, Netherlands) using a LiHa liquid handler (Tecan, Männedorf, Switzerland) to measure OD_660_ and external metabolites. 30 μL of each sample was diluted 5 fold in demineralized water in a 96 well plate and OD_660_ was measured with a Magellan Infinite 200 PRO spectrophotometer (Tecan, Männedorf, Switzerland). From the remaining sample, 150 μL was vacuum filter sterilized using 0.2 Multiscreen filter plates (Merck, Darmstadt, Germany) for HPLC measurements.

### 7-L wort fermentation cultivations

Batch cultivations under industrial conditions were performed in 10 L stirred stainless-steel fermenters containing 7 L of 16.6 °Plato wort. Fermentations were inoculated to a density of 5 × 10^6^ cells mL^−1^ at 8 °C. The temperature was raised during 48 hours to 11 °C and increased to 14 °C as soon as the gravity was reduced to 6.5 °Plato. Samples were taken daily during weekdays and the specific gravity and alcohol content were measured using an Anton Paar density meter (Anton Paar GmbH, Graz, Austria).

### Adaptive Laboratory Evolution

#### UV mutagenesis and selection

*S. eubayanus* CBS 12357^⊤^ was grown in a 500 mL shake flask containing 100 mL SMG at 20 °C until stationary phase and diluted to an OD_660_ of 1.0 with demineralized water. 50 mL of the resulting suspension was spun down at 4816 g for 5 min and washed twice with demineralized water. 25 mL of washed cells was poured into a 100 mm × 15 mm petri dish (Sigma-Aldrich) without lid and irradiated with a UV lamp (TUV 30 W T8, Philips, Eindhoven, The Netherlands) at a radiation peak of 253.7 nm. 25 mL of non-mutagenized and 5 mL of mutagenized cells were kept to determine survival rate. From both samples, a 100-fold dilution was made, from which successive 10-fold dilutions were made down to a 100,000-fold dilution. Then, 100 μL of each dilution was plated on YPD agar and the number of colonies were counted after incubation during 48h at room temperatures. After 10,000-fold dilution, 182 colonies formed from the non-mutagenized cells against 84 colonies for the mutagenized cells, indicating a survival rate of 46%. The remaining 20 mL of mutagenized cells was spun down at 4816 g for 5 min and resuspended in 1 mL demineralized water. Mutagenized cells were added to a 50 mL shake flask containing 9 mL SMMt and incubated for 21 days at 20 °C and 200 RPM. Maltotriose concentrations were analyzed at day 0, 19 and 21. After 21 days, two 100 μL samples were transferred to fresh shake flasks containing SMMt and incubated until stationary phase. At the end of the second transfer, single cell isolates were obtained using the BD FACSAria™ II SORP Cell Sorter (BD Biosciences, Franklin Lakes, NJ) equipped with a 488 nm laser and a 70 μm nozzle, and operated with filtered FACSFlow™ (BD Biosciences). Cytometer performance was evaluated by running a CST cycle with CS&T Beads (BD Biosciences). Drop delay for sorting was determined by running an Auto Drop Delay cycle with Accudrop Beads (BD Biosciences). Cell morphology was analysed by plotting forward scatter (FSC) against side scatter (SSC). Gated single cells were sorted into a 96 well microtiter plates containing SMMt using a “single cell” sorting mask, corresponding to a yield mask of 0, a purity mask of 32 and a phase mask of 16. The 96 well plates were incubated for 96 h at room temperature in a GENIos Pro micro plate spectrophotometer (Tecan, Männedorf, Switzerland), during which period growth was monitored as OD_660_. After 96 h, biomass in each well was resuspended using a sterile pin replicator and the final OD_660_ was measured. The 7 isolates with the highest final OD_660_ were picked, restreaked and stocked as isolates IMS0637-643. PCR amplification of the *S. eubayanus*-specific *SeFSY1* gene and ITS sequencing confirmed that all 7 isolates were *S. eubayanus*.

#### Laboratory evolution in chemostats

Chemostat cultivation was performed in Multifors 2 Mini Fermenters (INFORS HT, Velp, The Netherlands) equipped with a level sensor to maintain a constant working volume of 100 mL. The culture temperature was controlled at 20 °C and the dilution rate was set at 0.03 h^−1^ by controlling the medium inflow rate. Cultures were grown on 6-fold diluted wort supplemented with 10 g L^−1^ additional maltotriose (Glentham Life Sciences), 0.2 mL L^−1^ anti-foam emulsion C (Sigma-Aldrich, Zwijndrecht, the Netherlands), 10 mg L^−1^ ergosterol, 420 mg L^−1^ Tween 80 and 5 g L^−1^ ammonium sulfate. Tween 80 and ergosterol were added as a solution as described previously (61). IMS0637-IMS0643 were grown overnight at 20 °C and 200 RPM in separate shake flasks on 3-fold diluted wort. The OD_660_ of each strain was measured and the equivalent of 7 mL at an OD_660_ of 20 from each strain was pooled in a total volume of 50 mL. The reactor was inoculated by adding 20 mL of the pooled culture. After overnight growth, the medium inflow pumps were turned on and the fermenter was sparged with 20 mL min^−1^ of nitrogen gas and stirred at 500 RPM. The pH was not adjusted. Samples were taken weekly. Due to a technical failure on the 63^rd^ day, the chemostat was autoclaved, cleaned and restarted using a sample taken on the same day. After a total of 122 days, the chemostat was stopped and 10 single colony isolates were sorted onto SMMt agar using FACS, as for IMS0637-IMS0643. PCR amplification of the *S. eubayanus* specific *SeFSY1* gene and ITS sequencing confirmed that all ten single-cell isolates were *S. eubayanus*. Three colonies were randomly picked, restreaked and stocked as IMS0750-752.

#### Genomic isolation and whole genome sequencing

Yeast cultures were incubated in 50 mL Bio-One Cellstar Cellreactor tubes (Sigma-Aldrich) containing liquid YPD medium at 20°C on an orbital shaker set at 200 RPM until the strains reached stationary phase with an OD_660_ between 12 and 20. Genomic DNA for whole genome sequencing was isolated using the Qiagen 100/G kit (Qiagen, Hilden, Germany) according to the manufacturer’s instructions and quantified using a Qubit^®^ Fluorometer 2.0 (Thermo Scientific).

Genomic DNA of the strains CBS 12357^⊤^ and IMS0637-IMS0643 was sequenced by Novogene Bioinformatics Technology Co., Ltd (Yuen Long, Hong Kong) on a HiSeq2500 sequencer (Illumina, San Diego, CA) with 150 bp paired-end reads using PCR-free library preparation. Genomic DNA of the strains IMS0750 and IMS0752 was sequenced in house on a MiSeq sequencer (Illumina) with 300 bp paired-end reads using PCR-free library preparation. All reads are available at NCBI (https://www.ncbi.nlm.nih.gov/) under the bioproject accession number PRJNA492251.

Genomic DNA of strains IMS0637 and IMS0750 was sequenced on a Nanopore MinION (Oxford Nanopore Technologies, Oxford, United Kingdom). Libraries were prepared using 1D-ligation (SQK-LSK108) as described previously (80) and analysed on FLO-MIN106 (R9.4) flow cell connected to a MinION Mk1B unit (Oxford Nanopore Technology). MinKNOW software (version 1.5.12; Oxford Nanopore Technology) was used for quality control of active pores and for sequencing. Raw files generated by MinKNOW were base called using Albacore (version 1.1.0; Oxford Nanopore Technology). Reads with a minimum length of 1000 bp were extracted in fastq format. All reads are available at NCBI (https://www.ncbi.nlm.nih.gov/) under the bioproject accession number PRJNA492251.

#### Genome analysis

For the strains CBS 12357^⊤^, IMS0637-IMS0643, IMS0750 and IMS0752, the raw Illumina reads were aligned against a chromosome-level reference genome of CBS 12357^⊤^ (NCBI accession number PRJNA450912, https://www.ncbi.nlm.nih.gov/) (9) using the Burrows–Wheeler Alignment tool (BWA), and further processed using SAMtools and Pilon for variant calling (81-83). Heterozgous SNPs and INDELs which were heterozygous in CBS 12357^⊤^ were disregarded. Chromosomal translocations were detected using Breakdancer (84). Only translocations which were supported by at least 10% of the reads aligned at that locus were considered. Chromosomal copy number variation was estimated using Magnolya (85) with the gamma setting set to “none” and using the assembler ABySS (v 1.3.7) with a k-mer size of 29 (86). All SNPs, INDELs, recombinations and copy number changes were manually confirmed by visualising the generated .bam files in the Integrative Genomics Viewer (IGV) software (87). The complete list of identified mutations can be found in Supplementary Data File 1.

For strains IMS0637 and IMS0750, the nanopore sequencing reads were assembled de novo using Canu (version 1.3) (88) with –genomesize set to 12 Mbp. Assembly correctness was assessed using Pilon (83), and sequencing/assembly errors were polished by aligning Illumina reads with BWA (81) using correction of only SNPs and short indels (–fix bases parameter). Long sequencing reads of IMS0637 and IMS0750 were aligned to the obtained reference genomes and to the reference genome of CBS 12357^⊤^ using minimap2 (89). The genome assemblies for IMS0637 and IMS0750 are available at NCBI (https://www.ncbi.nlm.nih.gov/) under the bioproject accession number PRJNA492251.

#### Molecular biology methods

For colony PCR and Sanger sequencing, a suspension containing genomic DNA was prepared by boiling biomass from a colony in 10 μL 0.02 M NaOH for 5 min, and spinning cell debris down at 13,000 g. To verify isolates belonged to the *S. eubayanus* species, the presence of *S. eubayanus-* specific gene *SeFSY1* and the absence of *S. cerevisiae-specific* gene *ScMEX67* was tested by DreamTaq PCR (Thermo Scientific) amplification using primer pair 8572/8573 (90), and primer pair 8570/8571 (91), respectively. Samples were loaded on a 1% agarose gel containing SYBR Green DNA stain (Thermo Scientific). GeneRuler DNA Ladder Mix (Thermo Scientific) was used as ladder and gel was run at a constant 100V for 20 min. DNA bands were visualized using UV light. For additional confirmation of the *S. eubayanus* identity, ITS regions were amplified using Phusion High-Fidelity DNA polymerase (Thermo Scientific) and primer pair 10199/10202. The purified (GenElute PCR Cleanup Kit, Sigma-Aldrich) amplified fragments were Sanger sequenced (BaseClear, Leiden, Netherlands) (92). Resulting sequences were compared using BLAST to available ITS sequences of *Saccharomyces* species and classified as the species to which the amplified region had the highest sequence identity. The presence of the *SeMALT* genes was verified by using Phusion High-Fidelity DNA polymerase and gene specific primers: 10491/10492 for *SeMALT1*, 10632/10633 for *SeMALT2* and *SeMALT4/2*, 10671/10672 for *SeMALT3*, 10491/10671 for *SeMALT13*, and 10633/10671 for *SeMALT413*. The amplified fragments were purified using the GenElute PCR Cleanup Kit (Sigma-Aldrich) and Sanger sequenced (BaseClear) using the same primers used for amplification.

#### Plasmid construction

All plasmids and primers used in this study are listed in Table 3 and Supplementary Table S1, respectively. DNA amplification for plasmid and strain construction was performed using Phusion High-Fidelity DNA polymerase (Thermo Scientific) according to the supplier’s instructions. The coding region of *SeMALT413* was amplified from genomic DNA of IMS0750 with primer pair 10633/10671. Each primer carried a 40 bp extension complementary to the plasmid backbone of p426-TEF-amds (13), which was PCR amplified using primer pair 7812/5921. The transporter fragment and the p426-TEF-amdS backbone fragment were assembled (93) using NEBuilder HiFi DNA Assembly (New England Biolabs, Ipswich, MA), resulting in plasmid pUD814. The resulting pUD814 plasmid was verified by Sanger sequencing, which confirmed that its *SeMALT413* ORF was identical to the recombined ORF found in the nanopore assembly of IMS0750 (Figure 2C).

**Table 3:** Plasmids used during this study.

| Name | Relevant genotype | Source |
| --- | --- | --- |
| pUDP052 | *ori* (ColE1) *bla* panARSopt *amdSYM* *ScTDH3*<sub>pr</sub>-gRNA<sub>SeSGA1</sub>-*ScCYC1*<sub>ter</sub> *AaTEF1*<sub>pr</sub>-*Spcas9*<sup>D147Y P411T</sup>-*ScPHO5*<sub>ter</sub> | (9) |
| pUDE044 | *ori* (ColE1) *bla* 2μ *ScTDH3*<sub>pr</sub>-*ScMAL12*-*ScADH1*<sub>ter</sub> *URA3* | (94) |
| p426-TEF-amdS | *ori* (ColE1) *bla* 2μ *amdSYM* *ScTEF1*<sub>pr</sub>-*ScCYC1*<sub>ter</sub> | (13) |
| pUD479 | *ori* (ColE1) *bla* 2μ *amdSYM* *ScTEF1*<sub>pr</sub>-*SeMALT1*-*ScCYC1*<sub>ter</sub> | (9) |
| pUD480 | *ori* (ColE1) *bla* 2μ *amdSYM* *ScTEF1*<sub>pr</sub>-*SeMALT2*-*ScCYC1*<sub>ter</sub> | (9) |
| pUD814 | *ori* (ColE1) *bla* 2μ *amdSYM* *ScTEF1*<sub>pr</sub>-*SeMALT413*-*ScCYC1*<sub>ter</sub> | This study |

#### Strain construction

To integrate and overexpress *SeMALT2* and *SeMALT413* ORFs in *S. eubayanus* CBS 12357^⊤^, *SeMALT2* and *SeMALT413* were amplified from pUD480 and pUD814 respectively with primers 13559/13560 that carried a 40 bp region homologous to each flank of the *SeSGA1* gene located on *S. eubayanus* chromosome IX. To facilitate integration, the PCR fragments were co-transformed with the plasmid pUDP052 that expressed *Spcas9*^D147Y P411⊤^ (95, 96) and a gRNA targeting *SeSGA1* (9). The strain IMX1941 was constructed by transforming CBS 12357^⊤^ with 1 μg of the amplified *SeMALT2* expression cassette and 500 ng of plasmid pUDP052 by electroporation as described previously (96). Transformants were selected on SM_Ace_G plates. Similarly, IMX1942 was constructed by transforming CBS 12357^⊤^ with 1 μg of the amplified *SeMALT413* expression cassette for *SeMALT413* instead of *SeMALT2*. Correct integration was verified by diagnostic PCR with primer pair 12635/12636 (Supplementary Figure S8). All PCR-amplified gene sequences were Sanger sequenced (BaseClear).

#### Protein structure prediction

Homology modeling of the S*e*Malt413 transporter was performed using the SWISS-MODEL server (https://swissmodel.expasy.org/) (97). The translated amino acid sequence of *SeMALT413* was used as input (Supplementary Figure S3). The model of the xylose proton symporter XylE (PDB: 4GBY) was chosen as template (62). Models were built based on the target-template alignment using ProMod3. Coordinates which are conserved between the target and the template are copied from the template to the model. Insertions and deletions are remodeled using a fragment library. Side chains are then rebuilt. Finally, the geometry of the resulting model is regularized by using a force field. In case loop modelling with ProMod3 fails, an alternative model is built with PROMOD-II (98). 3D model was assessed and colored using Pymol (The PyMOL Molecular Graphics System, Version 2.1.1 Schrödinger, LLC.).

#### Sequence analysis of *SpMTY1*

The sequence of *SpMTY1* was analyzed by aligning *ScMAL31, ScAGT1, ScMPH2* and *ScMPH3* from *S. cerevisiae* strain S288C (63) and *SeMALT1, SeMALT2, SeMALT3, SeMALT4* from *S. eubayanus* strain CBS 12357^⊤^ (9) to the sequence of *SpMTY1* from *S. pastorianus* strain Weihenstephan 34/70 (22) using the Clone manager software (version 9.51, Sci-Ed Software, Denver, Colorado). The origin of nucleotides 969 to 1,639 of *SpMTY1* was further investigated using the blastn function of NCBI (https://www.ncbi.nlm.nih.gov/). The sequence was aligned against *S. cerevisiae* S288C (taxid:559292) to identify closely related homologues. In addition, *SpMTY1* was aligned against the complete nucleotide collection. To avoid similarity with genomes harboring an *MTY1* gene, sequences from *S. pastorianus* (taxid:27292), *S. cerevisiae* (taxid:4932), *S. eubayanus* (taxid:1080349), *S. cerevisiae* x *eubayanus* (taxid:1684324) and *S. bayanus* (*taxid:4931*) were excluded. The most significant alignment was with nucleotides 1,043,930 to 1,044,600 of chromosome VII of *S. paradoxus* strain YPS138 (GenBank: CP020282.1). As the most significant alignment of these nucleotides to *S. cerevisiae* S228C (taxid:559292) was *ScMAL31*, the gene was further referred to as *SparMAL31*.

### Analytics

The concentrations of ethanol and of the sugars glucose, maltose and maltotriose were measured using a high pressure liquid chromatography (HPLC) Agilent Infinity 1260 series (Agilent Technologies, Santa Clara, CA) using a Bio-Rad Aminex HPX-87H column at 65 °C and a mobile phase of 5 mM sulfuric acid with a flow rate of 0.8 mL per minute. Compounds were measured using a RID at 35 °C. Samples were spun down (13,000 g for 5 min) to collect supernatant or 0.2 μm filter-sterilized before analysis. The concentrations of ethylacetate and isoamylacetate, methanol, propanol, isobutanol, isoamyl alcohol and diacetyl were determined as described previously (60).

## Acknowledgments

We thank Jan-Maarten Geertman (Heineken Supply Chain B.V.) for his support during the study. This work was performed within the BE-Basic R&D Program (http://www.be-basic.org/), which was granted an FES subsidy from the Dutch Ministry of Economic Affairs, Agriculture and Innovation (EL&I).

## Supporting information captions

**Table S1:**
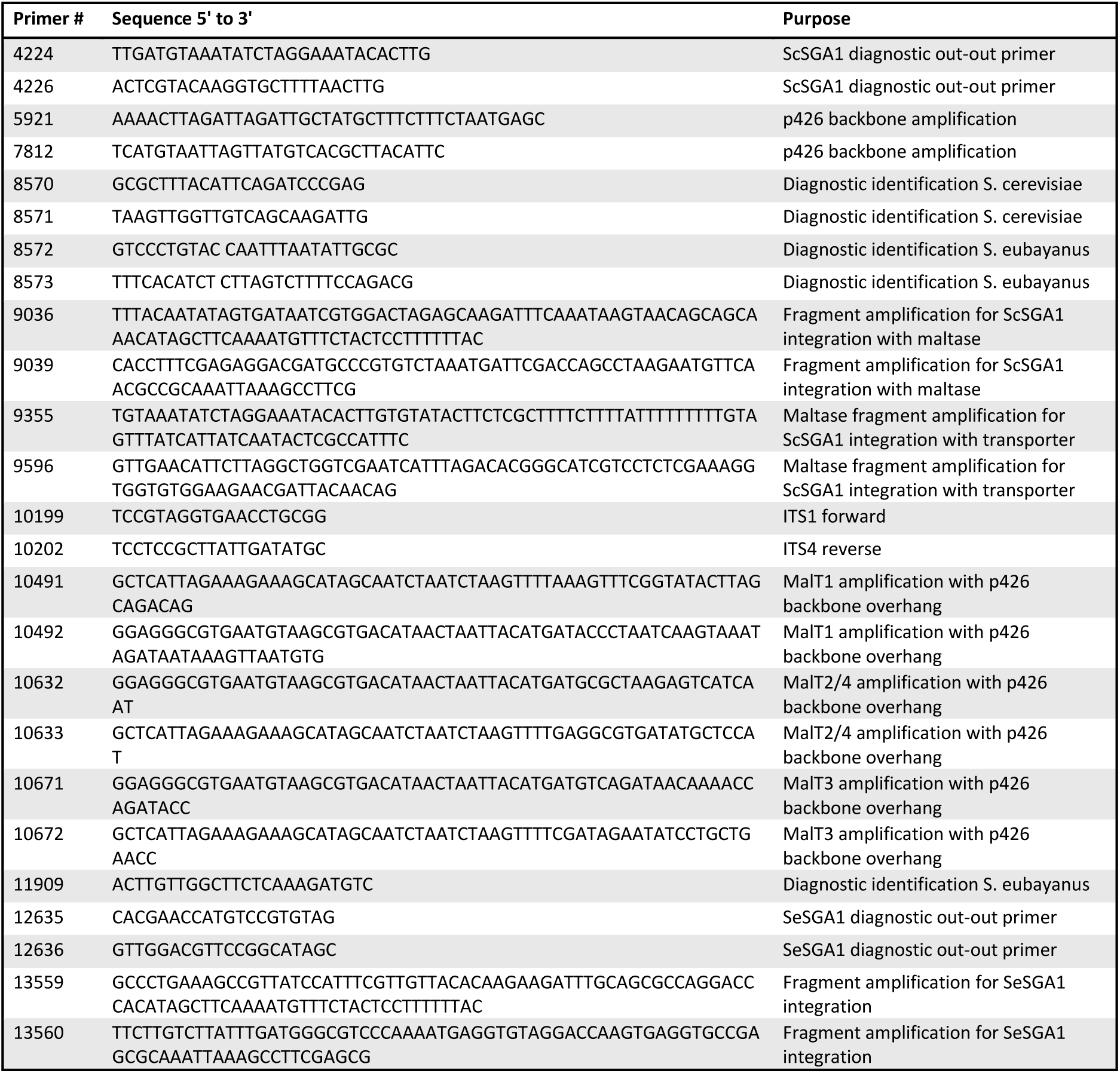
Primers used in this study.

**Figure S1:**
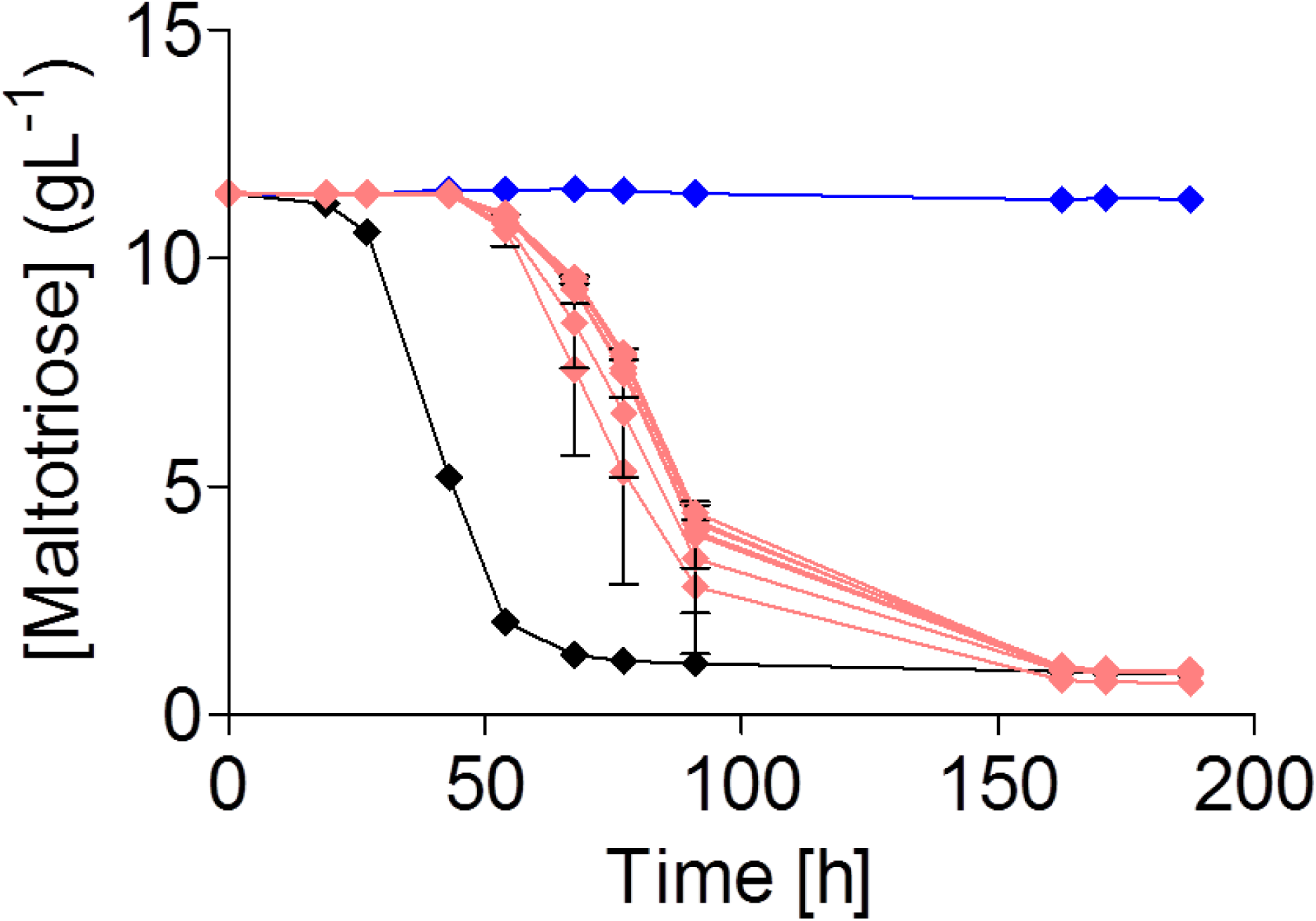
Characterization of enriched mutants IMS0637-IMS0643 on SMMt. Characterization of *S. pastorianus* CBS 1483 (black), *S. eubayanus* CBS 12357^⊤^ (blue) and selected mutants IMS0637-IMS0643 (light red) on SMMt at 20 °C. The average concentration of maltotriose (♦) and average deviation were determined from two replicates. IMS0637 was chosen as representative for all mutants.

**Figure S2:**
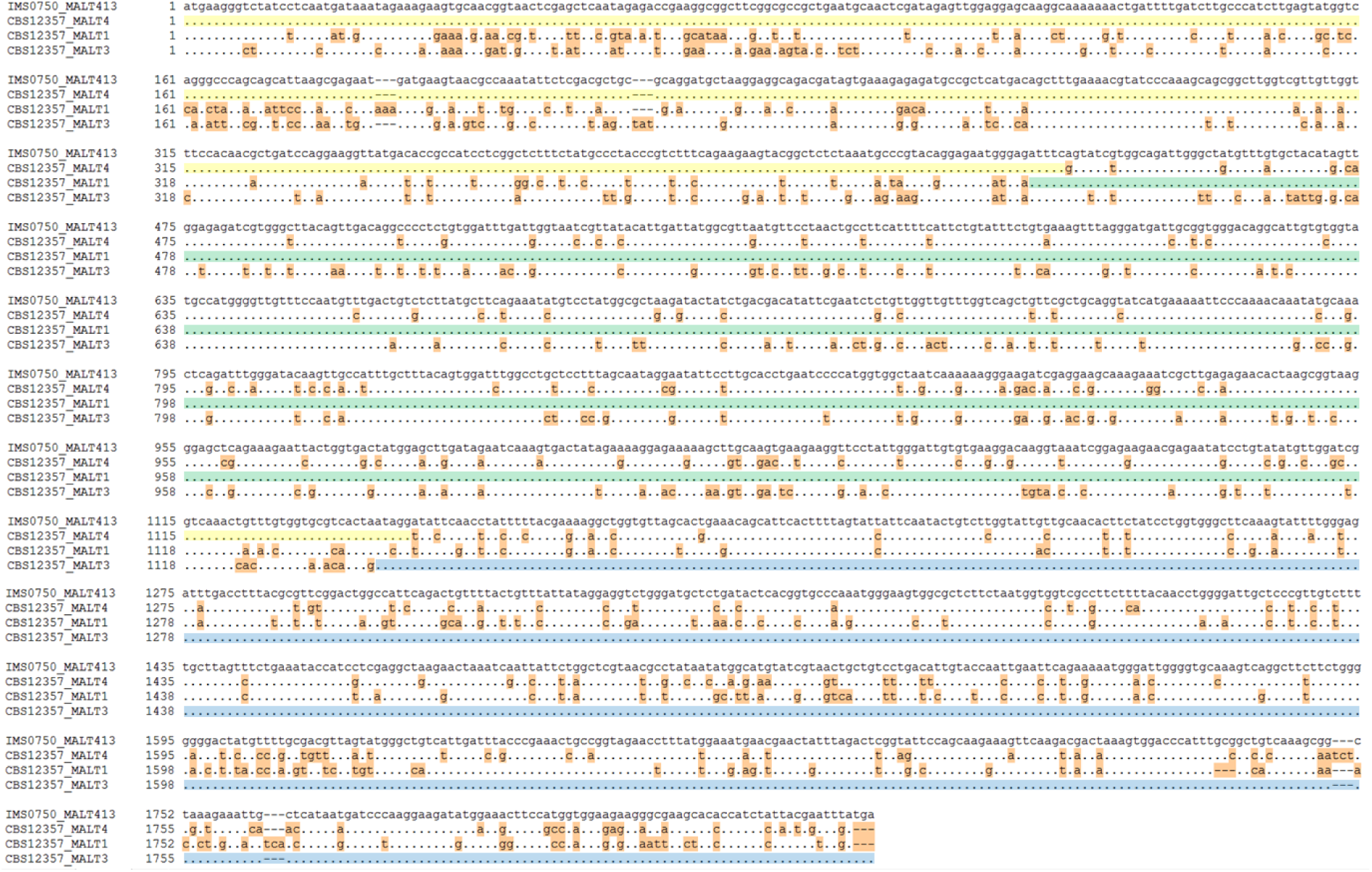
Sequence alignment of the *SeMALT* transporter genes from CBS 12357^⊤^ to the new recombined *SeMALT413* transporter gene from IMS0750. The allignment was performed using the Clone manager software (version 9.51, Sci-Ed Software). Identical nucleotides are shown by dots and nucleotides which differ from *SpMTY1* are shown in orange. In addition, the sequences which match exactly are highlighter in yellow for *SeMALT4*, in green for *SeMALT1* and in blue for *SeMALT3*.

**Figure S3:**
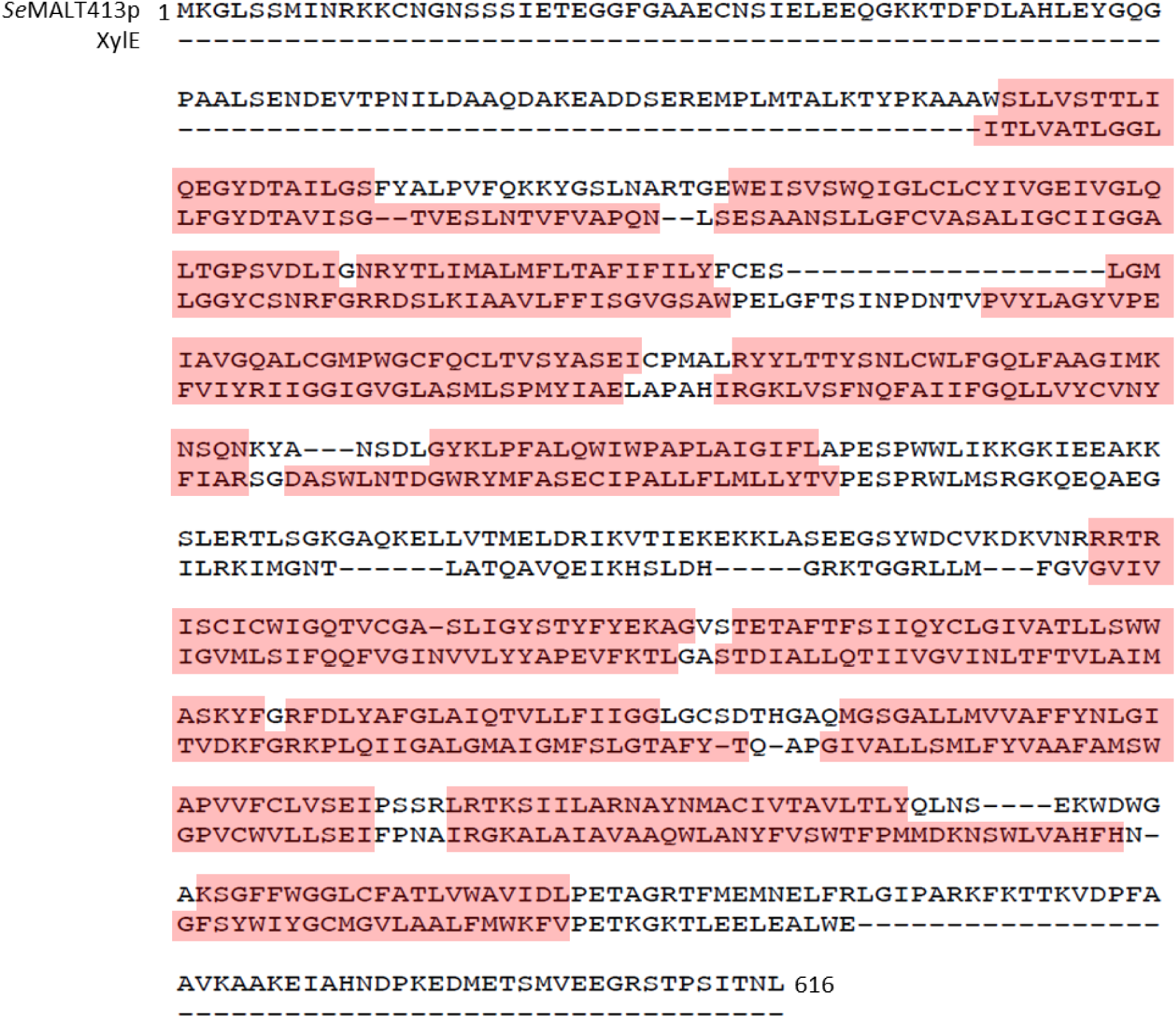
Alignment of SeMalt413 to XylE using by Promod3 for protein structure prediction. Transmembrane domain α-helices are indicated in red.

**Figure S4:**
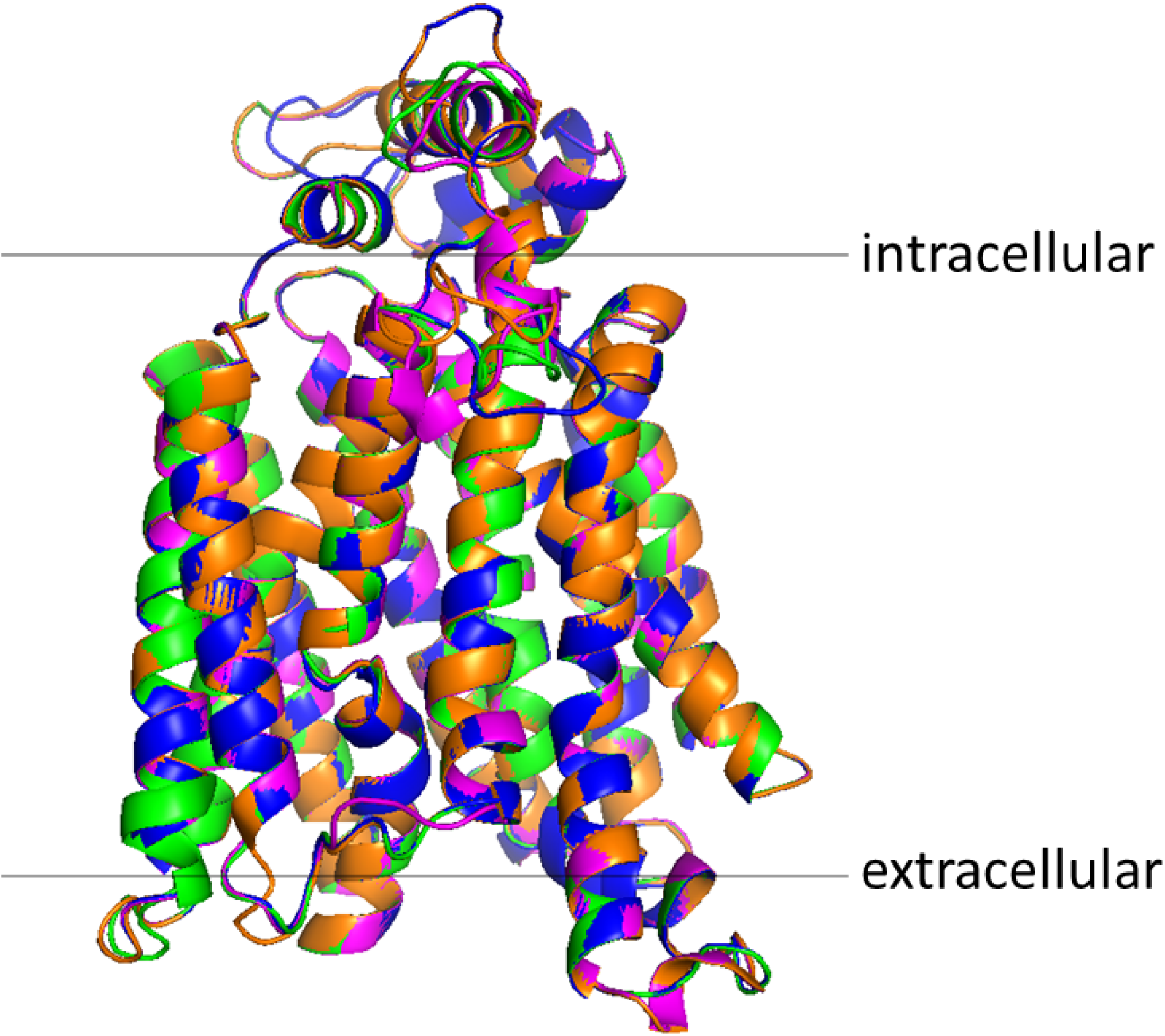
Protein structure overlay of S*e*Malt1 (green), S*e*Malt4 (orange), S*e*Malt3 (blue) and S*e*Malt413 (magenta). *SeMALT* genes were translated into amino acid sequence and used for structural prediction using SWISS-MODEL with XylE as a structural template. Resulting S*e*Malt protein structures were overlayed using PyMOL.

**Figure S5:**
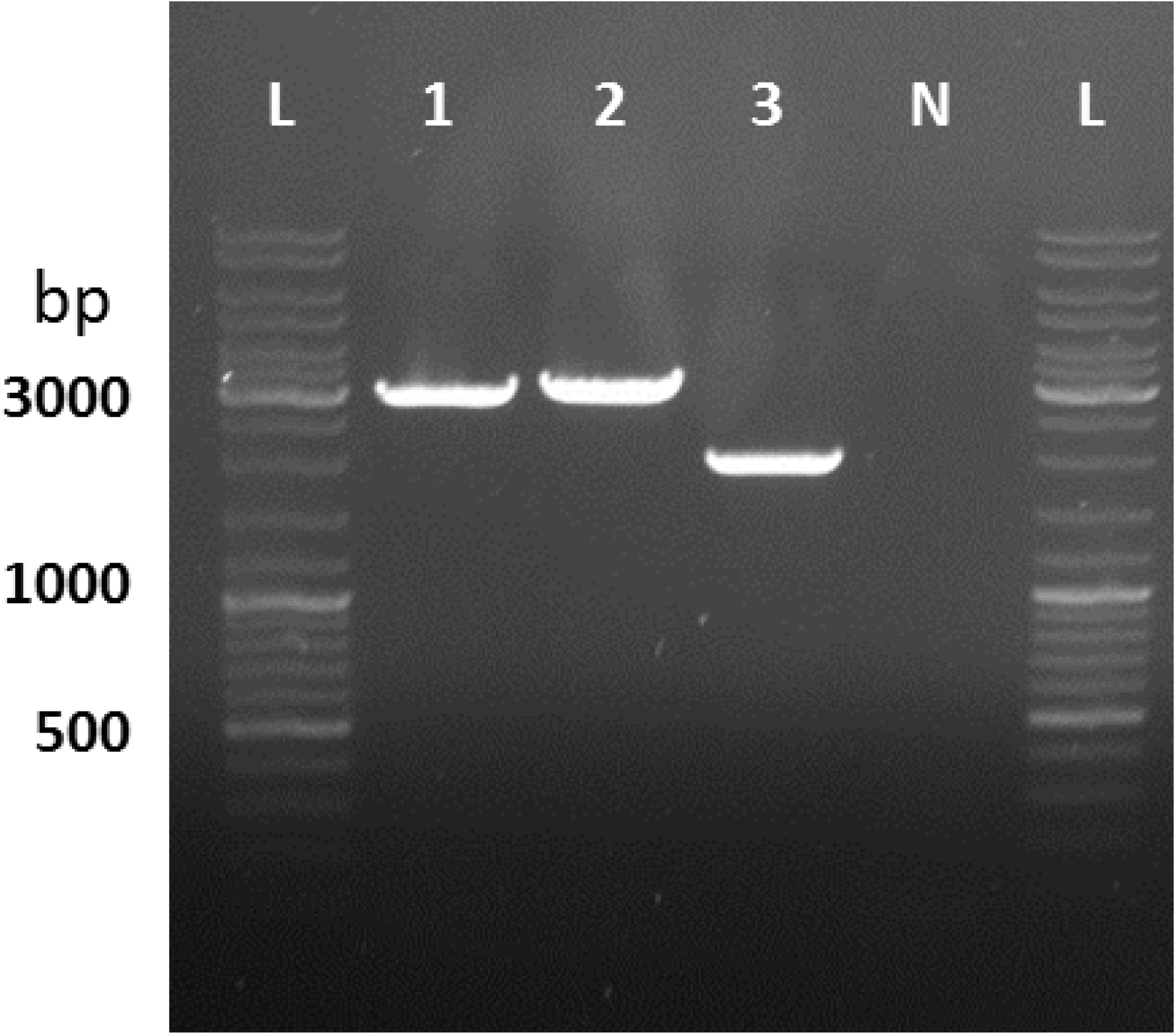
Amplification of the *SeSGA1* locus to verify integration of *SeMALT2* in IMX1941 and of *SeMALT413* in IMX1942. The *SeSGA1* locus was amplified from genomic DNA of IMX1941 (1), IMX1942 (2) and CBS 12357^⊤^ (3) using primers 12635/12636 and Phusion polymerase (Thermo Fischer Scientific). At the *SeSGA1* locus, IMX1941 should harbor *ScTEF1p-SeMALT2-ScCYC1t* and IMX1942 should harbor *ScTEF1p-SeMALT413-ScCYC1t*. As a negative control, a PCR was done with primers 12635/12636 without template DNA. L indicates the GeneRuler DNA Ladder Mix (Thermo Fischer Scientific).

**Figure S6:**
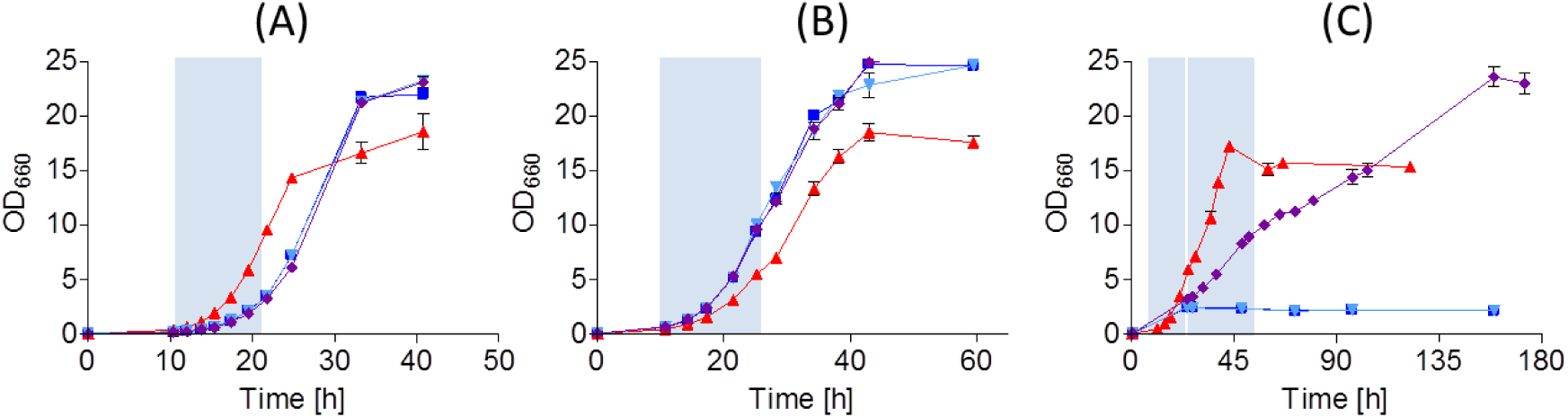
Characterization of (■) CBS 12357^⊤^, (▲) IMS0750, (▼) IMX1941, (♦) IMX1942 on SM (A) glucose, (B) maltose and (C) maltotriose. Strains were cultivated at 20 °C and optical densities were measured at 660 nm. Data represent average and standard deviation of three biological replicates. Blue boxes represent the timeframe used to calculate growth rates.

**Figure S7:**
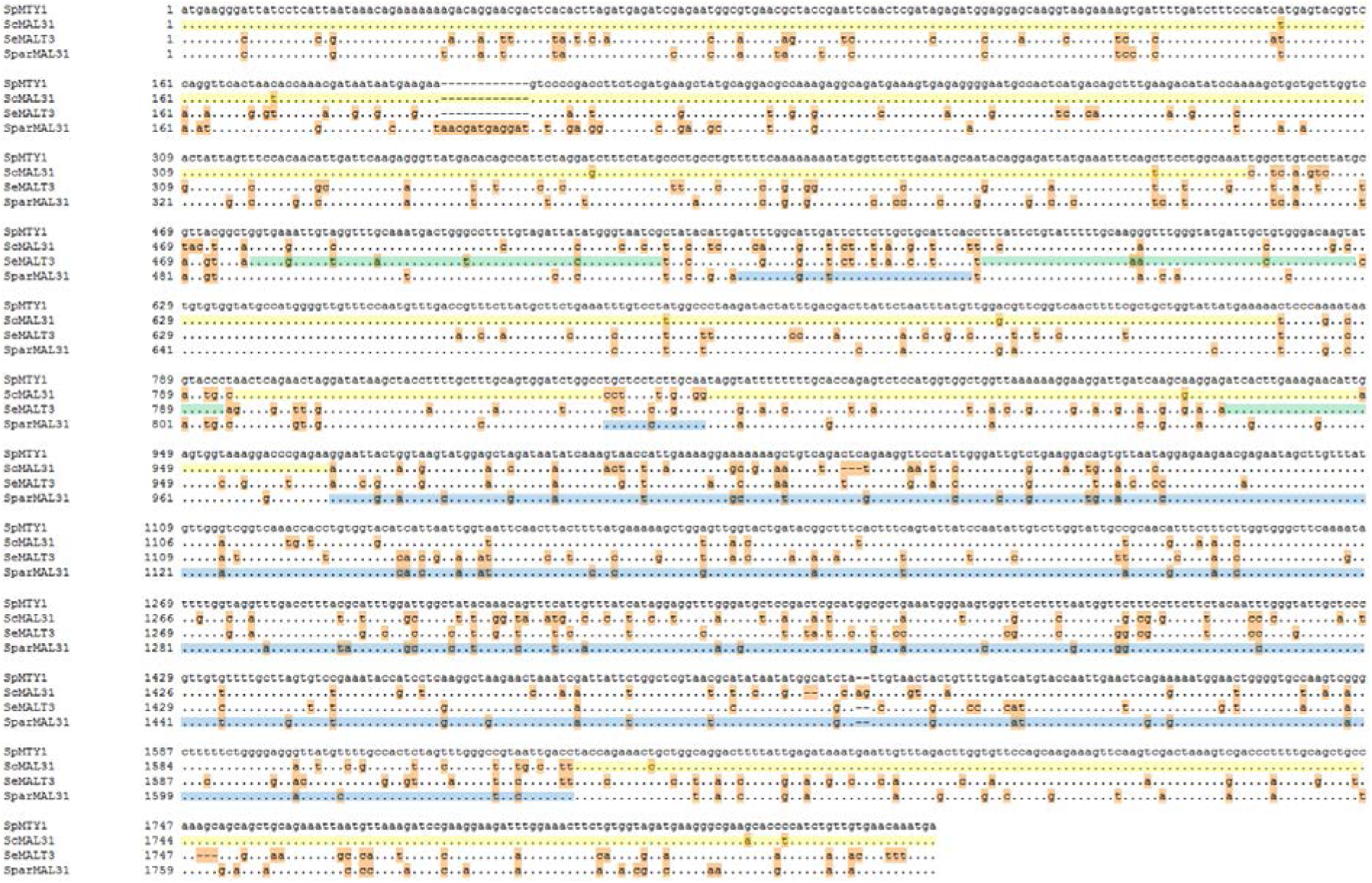
Alignment of *ScMAL31, SeMALT3* and *Spar*MAL31 to *SpMTY1* as obtained using Clone Manager. Identical nucleotides are shown by dots and nucleotides which differ from *SpMTY1* are shown in orange. In addition, the high-similarity sequences described in the main text are highlighted in yellow for *ScMAL31*, in green for *SeMALT3* and in blue for *SparMAL31*.

**Figure S8:**
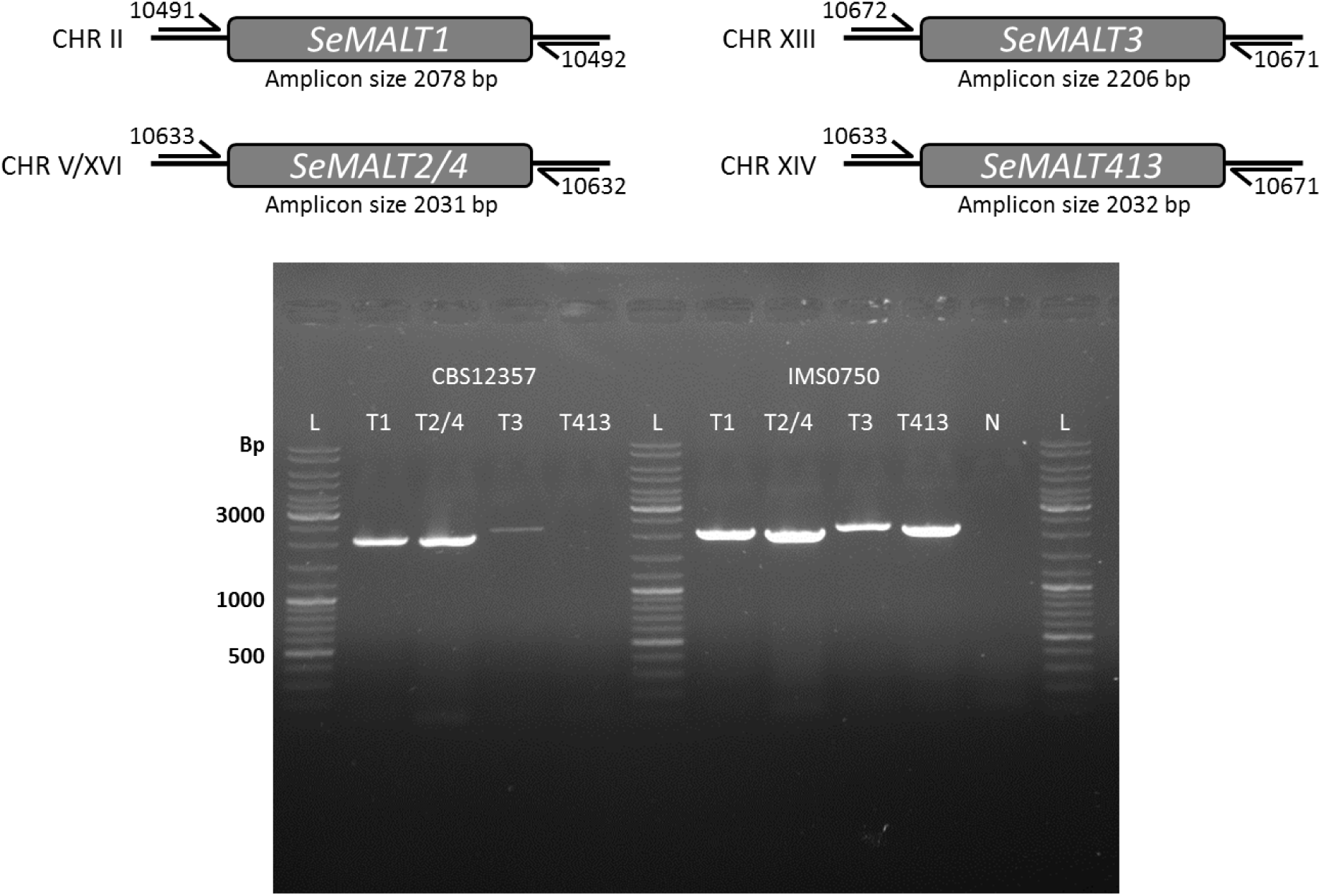
PCR amplification of *SeMALT* genes in wild type CBS 12357^⊤^ and evolved mutant IMS0750. The *SeMALT* genes were amplified from genomic DNA of CBS 12357^⊤^ and IMS0750 using Phusion polymerase (Thermo Fischer Scientific). Lanes show PCR products for *SeMALT1* (primers 10491/10492), *SeMALT2* and *SeMALT4* (primers 10633/10632), *SeMALT3* (primers 10672/10671) and *SeMALT413* (primers 10633/10671). As a negative control, a PCR was done with primers 10633/10632 without template DNA. L indicates the GeneRuler DNA Ladder Mix (Thermo Fischer Scientific).

